# The genetic architecture of *Arabidopsis thaliana* in response to native non-pathogenic leaf bacterial species revealed by GWA mapping in field conditions

**DOI:** 10.1101/2022.09.19.508615

**Authors:** Daniela Ramírez-Sánchez, Rémi Duflos, Chrystel Gibelin-Viala, Rémy Zamar, Fabienne Vailleau, Fabrice Roux

## Abstract

Non-pathogenic bacteria can largely contribute to plant health by mobilizing and supplying nutrients and by providing protection against pathogens and resistance to abiotic stresses. Yet, the number of GWAS reporting the genetic architecture of the response to individual members of the beneficial microbiota remains limited. In this study, we established a GWAS under field conditions to estimate the level of genetic variation and the underlying genetic architecture, among 162 accessions of *Arabidopsis thaliana* originating from 54 natural populations located south-west of France, in response to 13 strains of seven of the most abundant and prevalent non-pathogenic bacterial species isolated from the leaf compartment of *A. thaliana* in the same geographical region. Using a high-throughput phenotyping methodology to score vegetative growth-related traits, extensive genetic variation was detected within our local set of *A. thaliana* accessions in response to these leaf bacteria, both at the species and strain levels. The presence of crossing reaction norms among strains indicates that declaring a strain as a plant-growth promoting bacterium is highly dependent on the host genotype tested. In line with the strong genotype-by-genotype interactions, we detected a complex and highly flexible genetic architecture between the 13 strains. Finally, the candidate genes underlying the QTLs revealed a significant enrichment in several biological pathways, including cell, secondary metabolism, signalling and transport. Altogether, plant innate immunity appears as a significant source of natural genetic variation in plant-microbiota interactions and opens new avenues for better understanding the ecologically relevant molecular dialog during plant-microbiota interactions.

## INTRODUCTION

Both wild plant species and crops are consistently challenged by pathogens, making infectious disease often the major selective agent in nature [1–5]. In wild species, pathogen attacks can significantly decrease the number of offspring, which in turn affects host population growth rate [6–8]. Yield losses resulting from pathogen attacks can reach several tens of percent in crops [9–12], thereby threatening global food security [10, 13]. A major challenge in plant breeding and in ecological genomics is therefore to characterize the genetic architecture of response to pathogen attacks [14, 15]. Identifying the genetic and molecular bases for natural variation in response to pathogen attacks might lead to fundamental insights in the prediction of evolutionary trajectories of natural populations [16–19] and have enormous practical implications by increasing crop yield and quality [20–22].

Over the last decade, whole-genome sequencing made possible through the development of cutting-edge next-generation sequencing (NGS) technologies, combined with the development of increasingly sophisticated statistical methods in quantitative genetics [23, 24], led to a burst in the number of genome-wide association studies (GWAS) that were successful in both wild and cultivated plants. This allowed detecting genomic regions associated with natural variation of response to experimental inoculation with, in most cases, individual pathogenic strains [15, 25–27]. GWAS in plants revealed that the genetic architecture of response to pathogen attacks was highly polygenic [15], highly dependent on the abiotic environment [28, 29] and dynamic along the infection stages [28, 30]. In addition, the functional validation of few quantitative trait loci (QTLs) combined with transcriptomic analyses revealed both the involvement of a broad range of rarely considered molecular functions in plant immunity [17, 31, 32] as well as a new biomolecular network of the signaling machineries underlying disease resistance [33–35].

However, the entire set of microbial pathogens - also called pathobiota - represent only a small fraction of the entire set of microbes inhabiting plants, the so-called plant microbiota [14, 36, 37]. For instance, in the leaf compartment of 163 natural populations of *Arabidopsis thaliana* located in the south-west of France and characterized for bacterial communities using a metabarcoding approach allowing distinguishing pathogenic bacteria from other bacterial species [38], the relative abundance of pathobiota in microbiota was on average 1.6% in asymptomatic plants and 4.5% in plants with visible disease symptoms [38]. Furthermore, microbiota can largely contribute to plant health by (i) providing direct (production of antimicrobial components, niche competition) or indirect (triggering immune defense) protection against pathogens, (ii) mobilizing and provisioning nutrients, and (iii) providing resistance to abiotic stresses (such as drought) [37, 39–46]. Yet, the number of GWAS reporting the genetic architecture of the response to experimental inoculation with individual members of the beneficial microbiota remains limited in comparison to the number of GWAS on response to pathogens. In addition, despite the fact that the phyllosphere represents 60% of the total biomass on Earth and concentrating 10^26^ bacteria (Vorholt, 2012), most GWAS conducted in response to non-pathogenic bacteria focused on symbiotic bacteria or non-symbiotic plant-growth promoting bacteria (PGPB) at the below-ground level and in laboratory controlled conditions [47–52].

In this study, we established a GWAS under field conditions to estimate the level of genetic variation and the underlying genetic architecture, among 162 whole-genome sequenced accessions of *A. thaliana* originating from 54 natural populations located south-west of France, in response to 13 strains of seven of the most abundant and prevalent bacterial species isolated from the leaf compartment of *A. thaliana* in the same geographical region [53]. To do so, we first developed a high-throughput phenotyping methodology to score vegetative growth-related traits on tens of thousands of plants. We then combined GWA mapping derived from a Bayesian hierarchical model (BHM) [54], with a local score (LS) approach [55] to fine map QTLs down to the gene level, a combination that was successfully applied to detect and/or functionally validate QTLs involved in biotic interactions in *A. thaliana* [28, 30, 55, 56]. We finally identified the main biological pathways associated with all the candidate genes and discussed the function of the main candidate genes.

## MATERIAL AND METHODS

### Plant material

A total of 54 populations (each represented by three accessions) were chosen to represent both the genomic and ecological diversity identified among a set of 168 natural populations of *A. thaliana* located southwest of France [57, 58] (Supplementary Table S1). Seeds from maternal plants sampled in natural populations were collected in June 2016. Differences in the maternal effects among the 162 seed lots were reduced by growing one plant of each accession for one generation (Supplementary Text).

### Bacterial material

We considered two strains of seven (*i*.*e*. OTU2, OTU3, OTU4, OTU5, OTU6, OTU13 and OTU29) out of the 12 most abundant and prevalent non-pathogenic leaf bacterial OTUs identified across the 168 natural populations of *A. thaliana* [38], with the exception of OTU4 for which only one strain was available [53]. Based on whole-genome sequencing, the closest taxonomic classification for OTU2, OTU3, OTU4, OTU5, OTU6, OTU13 and OTU 29 was *Paraburkholderia fungorum, Oxalobacteraceae* bacterium, *Comamonadaceae* bacterium, *Pseudomonas moraviensis, Pseudomonas siliginis, Methylobacterium* sp. and *Sphingomonadaceae* bacterium, respectively [53]. For the purpose of another study, we also included the strain JACO-CL of the bacterial pathogen *Pseudomonas viridiflava* (OTU8), which is with *Xanthomonas campestris* the most abundant and prevalent bacterial pathogen across the 168 natural populations of *A. thaliana* [58].

### Experimental design and growth conditions

A field experiment of 15,552 plants was set up at the INRAE center of Auzeville-Tolosane using a split-plot design arranged as a randomized complete block design (RCBD) with 16 treatments nested within six experimental blocks (Figure S1). The 16 treatments correspond to two mock treatments and the individual inoculation of 14 bacterial strains, namely OTU2_*Pfu*_1, OTU2_*Pfu*_2, OTU3a_*Oxa*_1, OTU3a_*Oxa*_2, OTU4_*Com*_1, OTU5_*Pmo*_1, OTU5_*Pmo*_2, OTU6_*Psi*_1, OTU6_*Psi*_2, OTU13_*M*sp_1, OTU13_*M*sp_2, OTU29_*Sph*_1, OTU29_*Sph*_2 and OTU8_JACO-CL. Each block was represented by 48 trays of 54 individual bottom-pierced wells (Ø4.7 cm, vol. **∼**70 cm^2^) (SOPARCO, reference 4920) filled with PROVEEN® Semi-Bouturage 2. In each block, each treatment corresponded to three trays stuck to each other and containing 162 plants, with one replicate per accession (54 populations ^*^ 3 accessions). Randomization of accessions was kept identical among treatments within a block, but differed among the six blocks. Randomization of the 16 treatments differed between the six blocks, with the exception of the two mock treatments that were kept at the same position (Supplementary Figure S1).

All seeds were sown on March 18^th^ 2021, with several seeds sown in each well. Two weeks after sowing, seedlings were thinned to one per well, keeping the seedling the closest to the center of the well. During the entire growing period, the plants were watered as needed, *i*.*e*. manual watering morning and evening on hot and dry days and no watering on rainy days. A molluscicide (Algoflash® Naturasol) was regularly applied around the trays.

### Inoculation procedure

Bacterial strains were grown on solid medium in Petri dishes (TSA for OTUs 5, 6 and 8, TSB for OTU2, R2A for OTUs 3, 4 and 29, R2A for OTU13). The day of inoculation, bacterial colonies were resuspended in sterile deionized water and bacterial solutions were diluted to reach an OD_600 nm_ of 0.1. To facilitate the penetration of bacteria cells into plant organs, the Tween® 20 surfactant was added to each bacterial solution at a final concentration of 0.01%. Inoculation was performed 27 days after sowing (April 14, 2021), when most plants reached a 5-6 leaf stage. Using a Multipette® with a Combitips advanced® 50 mL, a volume of 1 mL of inoculum was dispensed on each rosette. A volume of 1mL of sterile water with a Tween® concentration of 0.01% was dispensed on each rosette of the plants of the two mock treatments. In order to increase relative humidity, plants were watered with a water mist spray system the seven days following the inoculation.

### Phenotyping

Following [59], a non-destructive imaging approach (Supplementary Figure S2) was used to measure each plant for nine traits related to vegetative growth (Supplementary Data Set 1): projected rosette surface area measured at 1day before inoculation (dbi) (**area-1dbi**), 5 days after inoculation (dai) (**area-5dai**) and 9 dai (**area-9dai**); rosette perimeter measured at 1 dbi (**perimeter-1dbi**), 5 dai (**perimeter-5dai**) and 9 dai (**perimeter-9dai**); maximal rosette diameter measured at 1 dbi (**diameter-1dbi**), 5 dai (**diameter-5dai**) and 9 dai (**diameter-9dai**). To estimate plant growth relative to size, three relative growth rates (RGR) were estimated based on the rosette surface area: RGR between 5 dai and 1 dbi (**RGR-5dai-1dbi**), RGR between 9 dai and 5 dai (**RGR-9dai-5dai**) and RGR between 9 dai and 1 dbi (**RGR-9dai-1dbi**). The procedure and methodologies are detailed in Supplementary Text.

### Data analyses

For the purpose of this study, the strain JACO-CL (OTU8, *P. viridiflava*) was not considered in any data analysis.

#### Investigation of the extent of natural genetic variation

To test the homogeneity of plant growth across the field trial and the presence of genetic variation for the three vegetative growth related traits measured before inoculation, data from the two mock treatments were pooled and the following mixed model (PROC MIXED procedure in SAS v. 9.4, SAS Institute Inc., Cary, NC, USA) was then used:

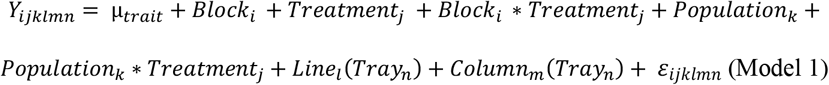

where Y is one of the three phenotypic traits measured before inoculation (*i*.*e*. area-1dbi, perimeter-1dbi and diameter-1dbi), μ is the overall mean of the phenotypic data, ‘Block’ accounts for differences in micro-environmental conditions among blocks, ‘Line(Tray)’ and ‘Column(Tray)’ accounts for difference in micro-environmental conditions within 54-well trays, ‘Treatment’ tests for difference among the 14 treatments (*i*.*e*. mock treatment and 13 treatments with non-pathogenic bacterial strains), ‘Population’ corresponds to the genetic differences among the 54 populations, ‘Population*Treatment’ tests whether the rank among the 54 populations differs among the 14 treatments, and ‘ε’ is the residual term.

While the terms ‘Treatment’ and ‘Population*Treatment’ were not significant, we detected a highly significant ‘Population’ effect (Supplementary Table S2), thereby indicating that the level of significant genetic variation observed among the 54 populations was homogeneous across the field trial before inoculation.

To estimate the natural genetic variation of the response of the 162 accessions nested within 54 populations to the 13 non-pathogenic bacterial strains, the following mixed model (PROC MIXED procedure in SAS v. 9.4, SAS Institute Inc., Cary, NC, USA) was used for each of the 15 treatments:

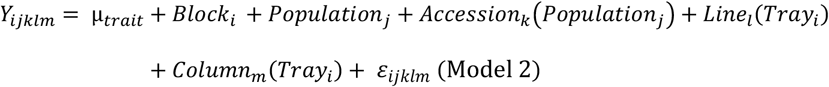

where Y corresponds to one of the nine traits (area-5dai, area-9dai, perimeter-5dai, perimeter-9dai, diameter-5dai, diameter-9dai, RGR-5dai-1dbi, RGR-9dai-5dai and RGR-9dai-1dbi). All the terms are identical to the ones described in Model (1), with the exception of ‘Accession’ that accounts for mean genetic differences among the three accession within populations.

For each of the 126 ‘phenotypic trait * treatment’ combinations (*i*.*e*. nine traits *14 treatments), genotypic values of the 54 populations were estimated by calculating least-squares (LS) mean values of the term ‘Population’ by the following linear model (PROC MIXED procedure in SAS v. 9.4, SAS Institute Inc., Cary, NC, USA) was used:

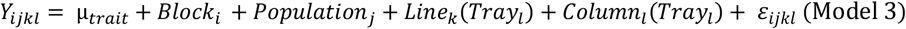

For each of the nine phenotypic traits, the estimated genotypic values (Supplementary Data Set 2) were then used to (i) compare phenotypic variation among the 14 treatments, (ii) estimate the level of ‘Population*Treatment’ interactions by calculating pairwise non-linear correlation coefficients (Spearman’s *rho*) among the 14 treatments, and (iii) run GWA analyses (see below).

To estimate broad-sense heritability values (*H*^2^) for each of the 126 ‘phenotypic trait * treatment’ combinations, the following linear model (PROC MIXED procedure in SAS v. 9.4, SAS Institute Inc., Cary, NC, USA) was used:

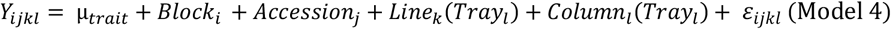

After considering the effects of the terms ‘Line(Tray)’ and ‘Column(Tray)’, the percentage of phenotypic variance explained by each other term of Model 4 was estimated by the PROC VARCOMP procedure (REML method, SAS v. 9.4, SAS Institute Inc., Cary, NC, USA). Following [33], *H*^2^ values were estimated using the following formula:

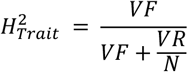

where ‘VF’ corresponds to the genetic variance among the 162 accessions, “VR” is the residual variance, and ‘*N*’ is the mean number of biological replicates per accession (N = 6 in this study). In Models 1, 2, 3 and 4, all factors were treated as fixed effects. For calculating *F*-values, terms were tested over their appropriate denominators. A correction for the number of tests was performed to control the False Discover Rate (FDR) at a nominal level of 5%.

#### Combining GWA mapping with a local score approach (GW-LS)

Based on a Pool-Seq approach, a representative picture of within-population genetic variation was previously obtained for 168 natural populations of *A. thaliana* located southwest of France [58], leading to the estimation of standardized allele frequencies corrected for the effect of population structure within each population for 1,638,649 SNPs across the genome [57, 58]. For the purpose of this study, standardized population allele frequencies were retrieved for the 54 populations. Then, for each of the 126 ‘phenotypic trait * treatment’ combinations, a genome scan was first launched by estimating for each SNP Spearman’s *rho* and associated *p* values between standardized allele frequencies and population genotypic values. Thereafter, to increase (i) the resolution in fine mapping genomic regions associated with genetic variation in response to bacterial strains, and (ii) the identification of QTLs with small effects, we followed [28, 30, 55, 56] by implementing a local score approach (with tuning parameter ξ = 2) on these *p* values. Finally, significant SNP-phenotype associations were identified by estimating a chromosome-wide significance threshold for each chromosome [55].

#### Enrichment in biological processes

A custom script written under the *R* environment [56] was used to retrieve the candidate genes underlying detected QTLs for each of the 126 ‘phenotypic trait * treatment’ combinations. For each of the 14 treatments, we merged the lists of candidate genes of the nine phenotypic traits and removed duplicates. For each of the 13 treatments with a non-pathogenic bacterial strain, only candidate genes not found in the mock treatment were kept. To identify biological pathways significantly over-represented (*P* < 0.01), each of the 14 resulting lists of unique candidate genes were submitted to the classification SuperViewer tool on the university of Toronto website (http://bar.utoronto.ca/ntools/cgibin/ntools_classification_superviewer.cgi) using the MAPMAN classification.

## RESULTS

### Genetic variation of *Arabidopsis thaliana* in response to non-pathogenic bacterial strains in field conditions

In agreement with previous experiments conducted in *in vitro* conditions [53], no disease symptoms were observed in our field conditions. For each of the 14 treatments (mock treatment and 13 treatments with a non-pathogenic bacterial strain), highly significant genetic variation was detected both between the 54 populations (Figure 1, Supplementary Figure S3, Supplementary Table S3) and between the 162 accessions (Supplementary Table S4) for each of the nine phenotypic traits, with the exception of (i) the rosette perimeter at 9 dai in presence of OTU3a_*Oxa*_1 at the population level (Supplementary Table S3), and (ii) the rosette area at 9 dai in presence of OTU5_*Pmo*_1 and OTU6_*Psi*_1 at the accession level (Supplementary Table S4). Across the 126 ‘phenotypic trait * treatment’ combinations, the mean broad-sense heritability (*H*^2^) estimate was 0.49 (median = 0.56, quantile 5% = 0.16, quantile 95% = 0.70). These results indicate that a non-negligible fraction of phenotypic variance was explained by genetic variation among populations and accessions (Supplementary Table S4).

**Figure 1.**
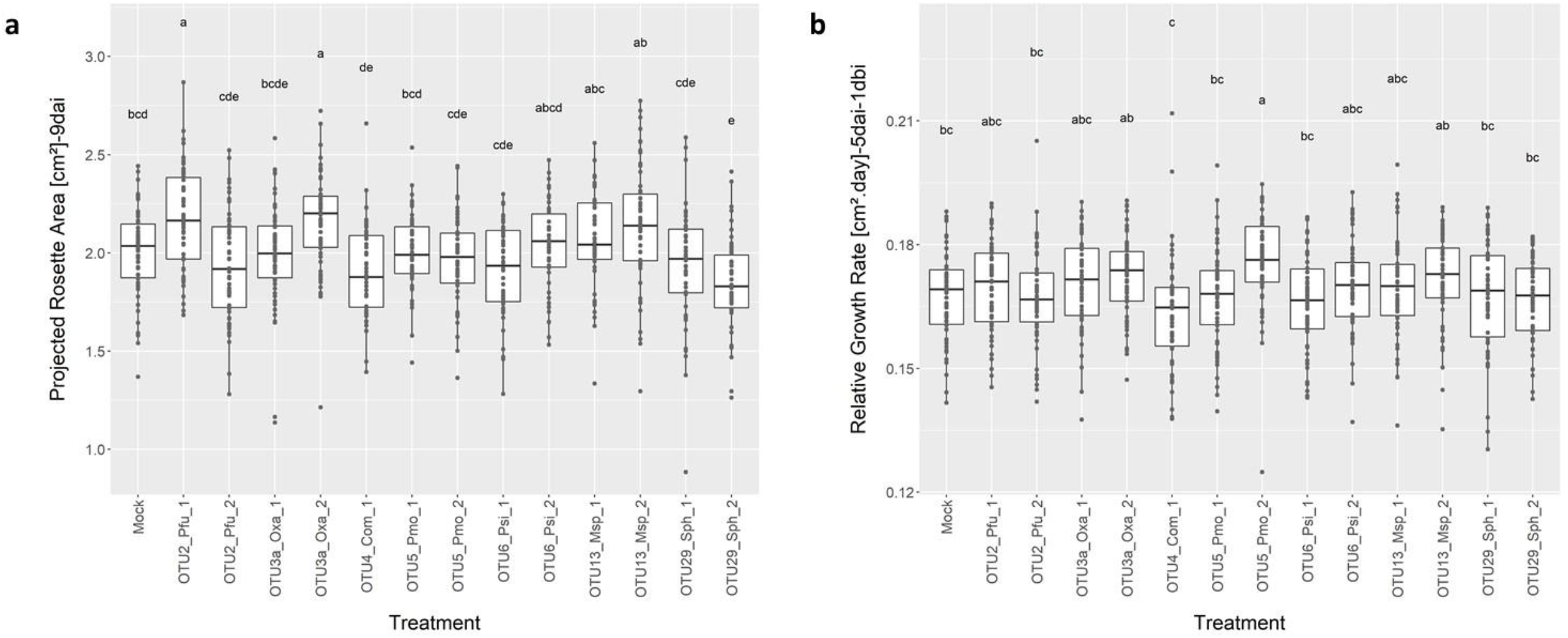
Phenotypic variation of the response to the mock treatment and the 13 bacterial strains in field conditions. **a** Box-plots illustrating the variation among the 14 treatments for the trait ‘area-9dai’. **b** Box-plots illustrating the variation among the 14 treatments for the trait ‘RGR-5dai-1dbi’. For each treatment, each dot corresponds to the genotypic value of one of the 54 populations of *A. thaliana*. For each trait, different letters indicate different groups according to the treatments after a Ryan-Einot-Gabriel-Welsh (REGWQ) multiple-range test at *P* = 0.05. dai: days after inoculation, dbi: day before inoculation.

A significant variation was observed among the 14 treatments for each phenotypic trait (Figure 1, Supplementary Figure S3). However, significant differences between the response to any bacterial strain and the mock treatment were only observed for three traits (*i*.*e*. area-9dai, diameter-9dai and RGR-5dai-1dbi) (Figure 1, Supplementary Figure S3). For instance, the rosette area at 9 dai was on average bigger and smaller in response to OTU2_*Pfu*_1/OTU3a_*Oxa*_2 and OTU29_*Sph*_2 than in the mock treatment, respectively (Figure 1a). The relative growth rate between 5 dai and 1 dbi was significantly higher in response to OTU5_*Pmo*_2 than in seven treatments, including the mock treatment (Figure 1b). More importantly, for each phenotypic trait, we observed a strong genetic variation among the 54 populations in their response to each of the 13 non-pathogenic bacterial strains (Figure 2, Supplementary Figure 4). Indeed, values of genetic correlations between the mock treatment and each treatment with a bacterial strain were largely deviating from 1, in particular at 9 dai (Figure 2a), with the exception of relative growth rate for with lower values of genetic correlations were observed within 5 dai than within 9 dai (Figure 2b, Supplementary Figure S4). In addition, the response of the 54 populations greatly varied among the 13 bacterial strains, even between two strains belonging to the same bacterial species (Figure 2, Supplementary Figure 4). For instance, while most populations present either a positive, neutral or negative response to either OTU13_*M*sp strain (*i*.*e*. presence of crossing reaction norms), the direction and/or the strength of response of given population can largely differ between the two OTU13_*M*sp strains, as illustrated by the populations FERR-A and LUZE-B (Figure 3).

**Figure 2.**
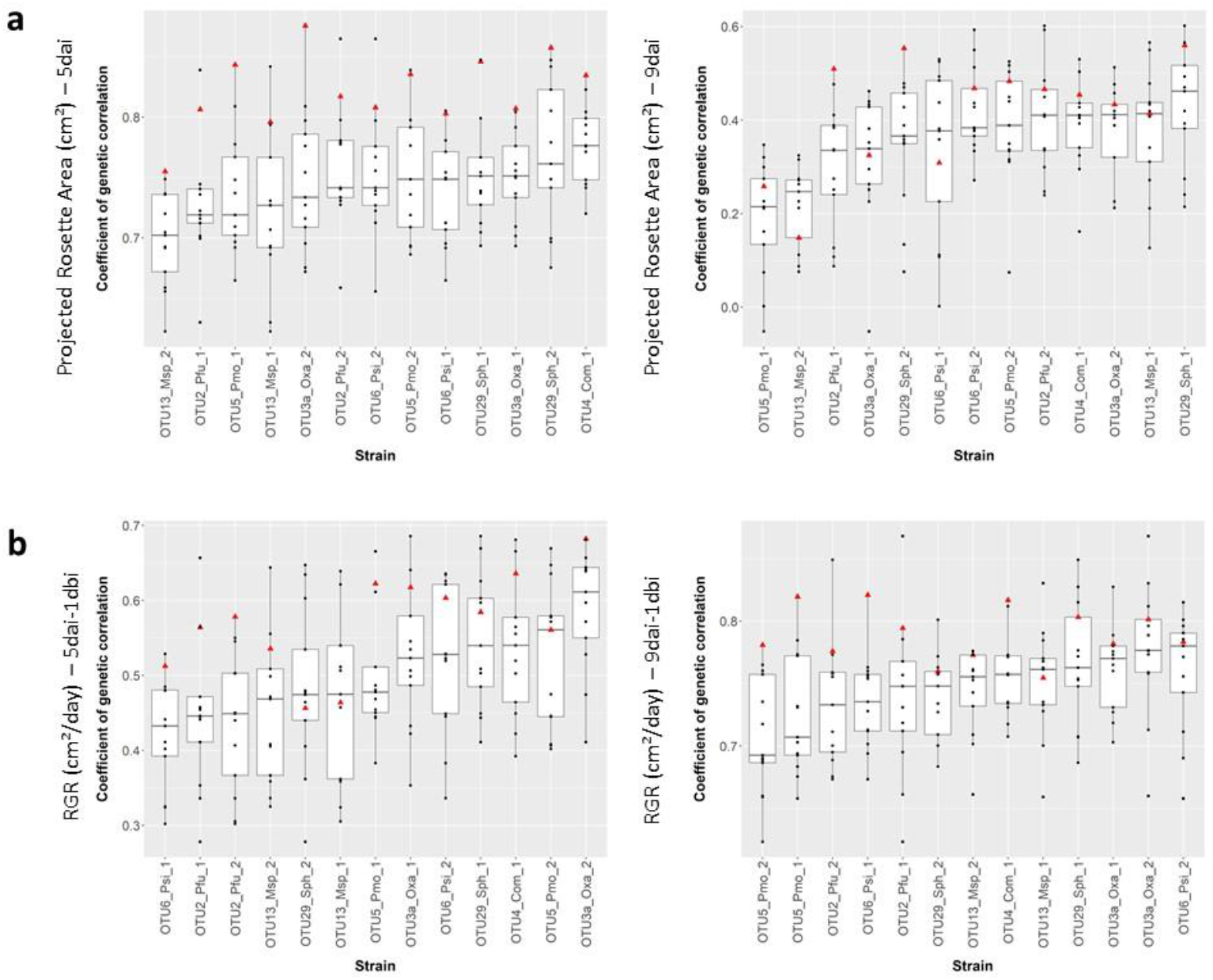
Genetic variation of 54 natural populations of *A. thaliana* in response to the 13 bacterial strains in field conditions. **a** Box-plots illustrating the range of genetic correlations between each treatment with a bacterial strain and the remaining 13 treatments for the traits ‘area-5dai’ and ‘area-9dai’. **b** Box-plots illustrating the range of genetic correlations between each treatment with a bacterial strain and the remaining 13 treatments for the traits ‘RGR-5dai-1dbi’ and ‘RGR-9dai-1dbi’. Red triangle: genetic correlation with the mock treatment, black dots: genetic correlations with other treatments with a bacterial strain (*ggplot2* library implemented in the R environment). dai: days after inoculation, dbi: day before inoculation. Treatments are ranked according to their mean genetic correlation with other treatments.

**Figure 3.**
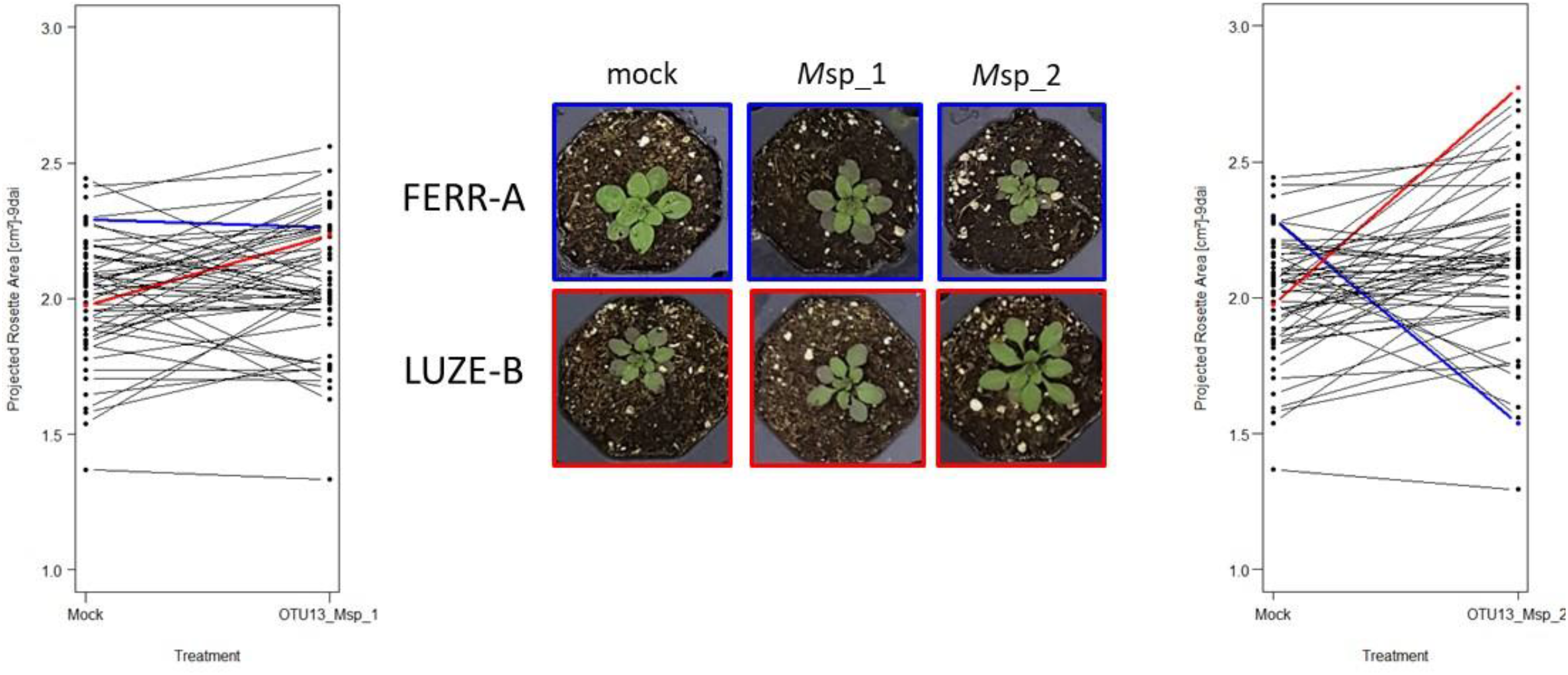
Interaction plots illustrating the reaction norms observed at the population level between the mock treatment and the treatment with OTU13_*M*sp_1 (left panel) and OTU13_*M*sp_2 (right panel). Each dot corresponds to the genotypic value of one of the 54 populations of *A. thaliana*. Each line corresponds to the response of one of the 54 populations to the inoculation with either OTU13_*M*sp strain. The blue and red lines correspond to two populations FERR-A and LUZE-B, respectively, with an opposite response to the strain OTU13_*M*sp_2. Pictures illustrate representative plants of the two populations highlighted in blue and red for the mock treatment and the treatment with either OTU13_*M*sp strain.

Altogether, the presence of (i) genetic variation at the population and accession levels for most ‘phenotypic trait * treatment’ combinations, (ii) crossing reaction norms between the mock treatment and each treatment with a bacterial strain, and (ii) crossing reaction norms among the 13 treatments with a bacterial strain, suggests a genetic architecture that largely differs among the 14 treatments, whatever the phenotypic trait considered.

### A genomic map of local adaptation to prevalent and/or abundant leaf bacterial species

Based on the allele frequencies of 1,638,649 SNPs obtained by a Pool-Seq approach for each of the 54 populations (Frachon et al., 2018), a GWA mapping analysis combining a Bayesian hierarchical model with a local score approach (BMH-LS) was conducted to characterize the genetic architecture of response to the 13 non-pathogenic bacterial strains. Across the 126 ‘phenotypic trait * treatment’ combinations, we detected 2,064 QTLs with a mean length of QTL interval equal to **∼**837bp (quantile 5% **∼** 38bp, quantile 95% = 3.12kb) (Supplementary Data Set 3). The number of QTLs per ‘phenotypic trait * treatment’ combination ranged from 6 to 34 (mean =16.4), suggesting a polygenic architecture for the response to members of the most prevalent and/or abundant non-pathogenic bacterial species of the leaf compartment of *A. thaliana* located south-west of France (Figure 4a).

**Figure 4.**
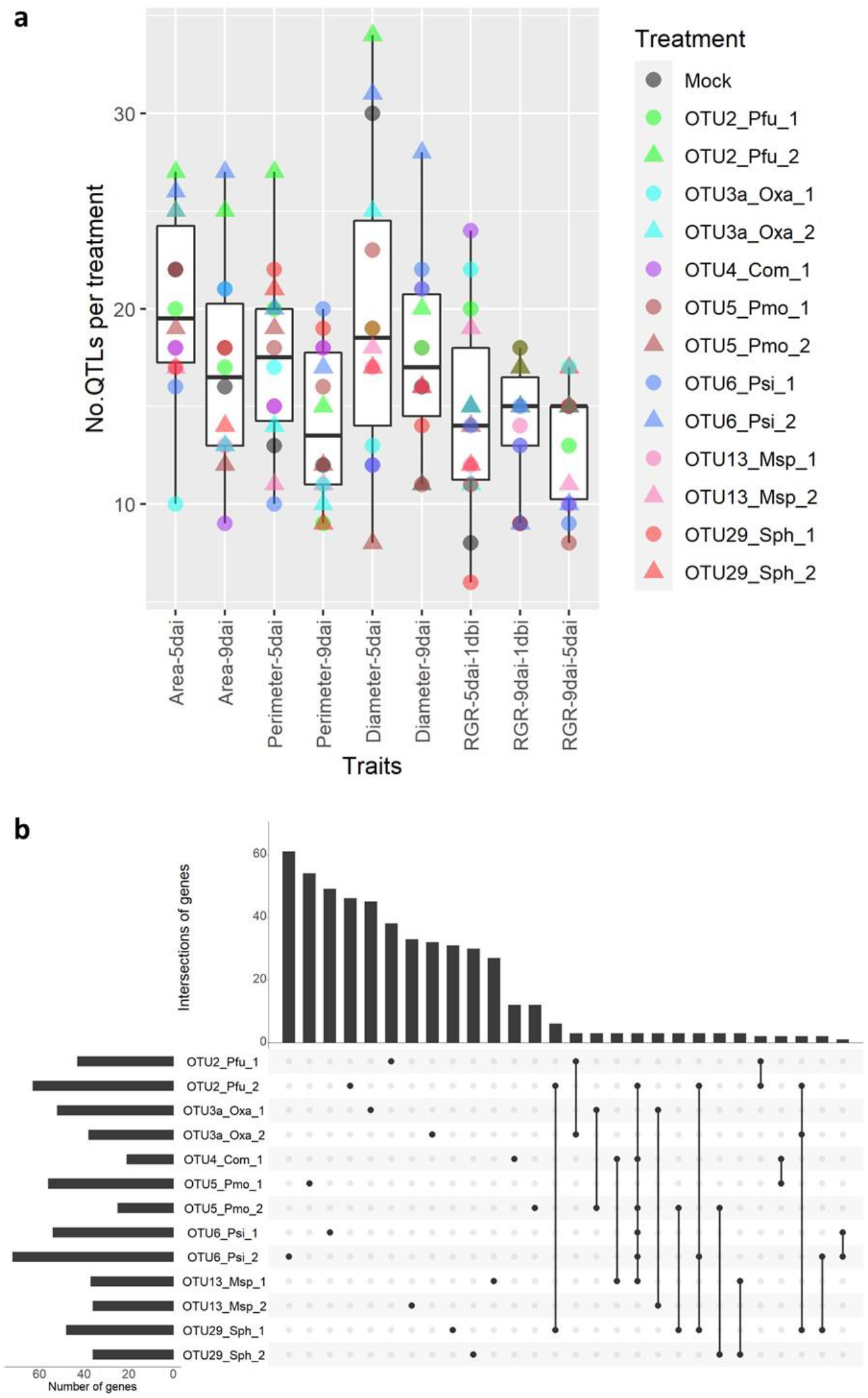
Genetic architecture of the response of 54 natural populations of *A. thaliana* to the 13 bacterial strains in field conditions. **a** Number of QTLs per treatment for each of the nine phenotypic traits. **b** An UpSet plot illustrating the flexibility of genetic architecture among the 13 treatments with bacterial strains for the trait ‘area-9dai’ (*upset* library implemented in the *R* environment). ‘Number of genes’: Total number of candidate genes underlying detected QTLs and not shared with the mock treatment. A single dot indicates the number of candidate genes specific to a given treatment. Candidate genes shared between two or more treatments are represented by a line connecting two or more dots.

In agreement with the level of genetic correlations observed among the 14 treatments (mock treatment and 13 treatments with a non-pathogenic bacterial strain) and the presence of crossing reaction norms (Figures 2 and3, Supplementary Figures S3 and S4), the genetic architecture was highly flexible between the mock treatment and treatments with bacterial strains, as well as among treatments with bacterial strains at the interspecific and intraspecific levels, as illustrated for the rosette area at 9 dai (Figure 5). For instance, most candidate genes underlying detected QTLs and not shared with the mock treatment were specific to a given treatment with a bacterial strain (Figure 4b, Supplementary Figure 5, Supplementary Data Set 4), in particular at 9 dai (Figure 4b, Supplementary Figure 5). For instance, for the maximal rosette diameter, while the percentage of candidate genes specific to a given treatment with a bacterial strain ranged from 57.7% to 86.1% (mean = 75.2%) at 9 dai, it ranged from 26.9% to 81.1% (mean = 46.0%) at 5 dai (Supplementary Figure 5).

**Figure 5.**
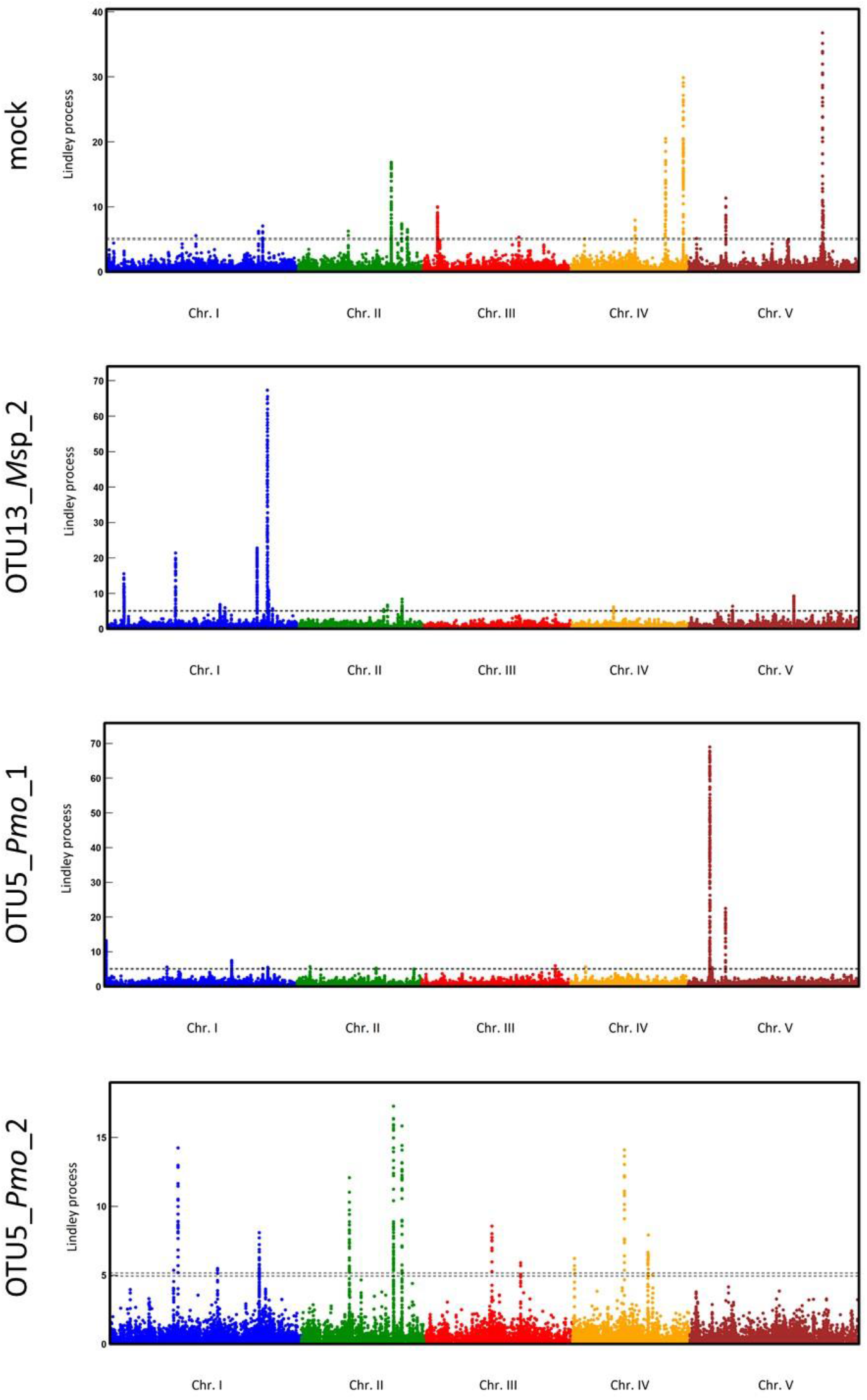
Manhattan plots of the Lindley process for the trait ‘area_9dai’ for the mock treatment and the treatments with the bacterial strains OTU13_*M*sp_2, OTU5_*Pmo*_1 and OTU5_*Pmo*_2. The *x*-axis corresponds to the physical position of 1,638,649 SNPs on the five chromosomes. The dashed lines indicate the minimum and maximum of the five chomosome-wide significance thresholds.

### Identification of enriched biological processes and candidate genes associated with the response to prevalent and/or abundant leaf bacterial species

The first approach to identify relevant candidate genes involved in the response to the 13 non-pathogenic bacterial species was to focus on candidate genes underlying the most pleiotropic QTLs. Here, we focused on QTLs detected for the response in more than six bacterial strains, but not detected for the mock treatment. We identified seven such pleiotropic QTLs encompassing 17 candidate genes (Table 1, Supplementary Data Set 5). In agreement with the highly flexible genetic architecture observed between strains within a bacterial species (Figure 4b, Supplementary Figure 5), the high level of pleiotropy observed for these QTLs was more dependent on the identity of the bacterial strains than the identity of the bacterial species (Table 1). Among the 17 candidate genes, eight genes have functions in relation with plant development and organ growth, *i*.*e. At2g40650* [60], *At2g40670* [61, 62], *At2g44710* [63], *At2g47190* [64–66], *At4g14713* [67–70], *At4g14716* [71], *At4g14720* [72] and *At5g42360* [73].

**Table 1.**
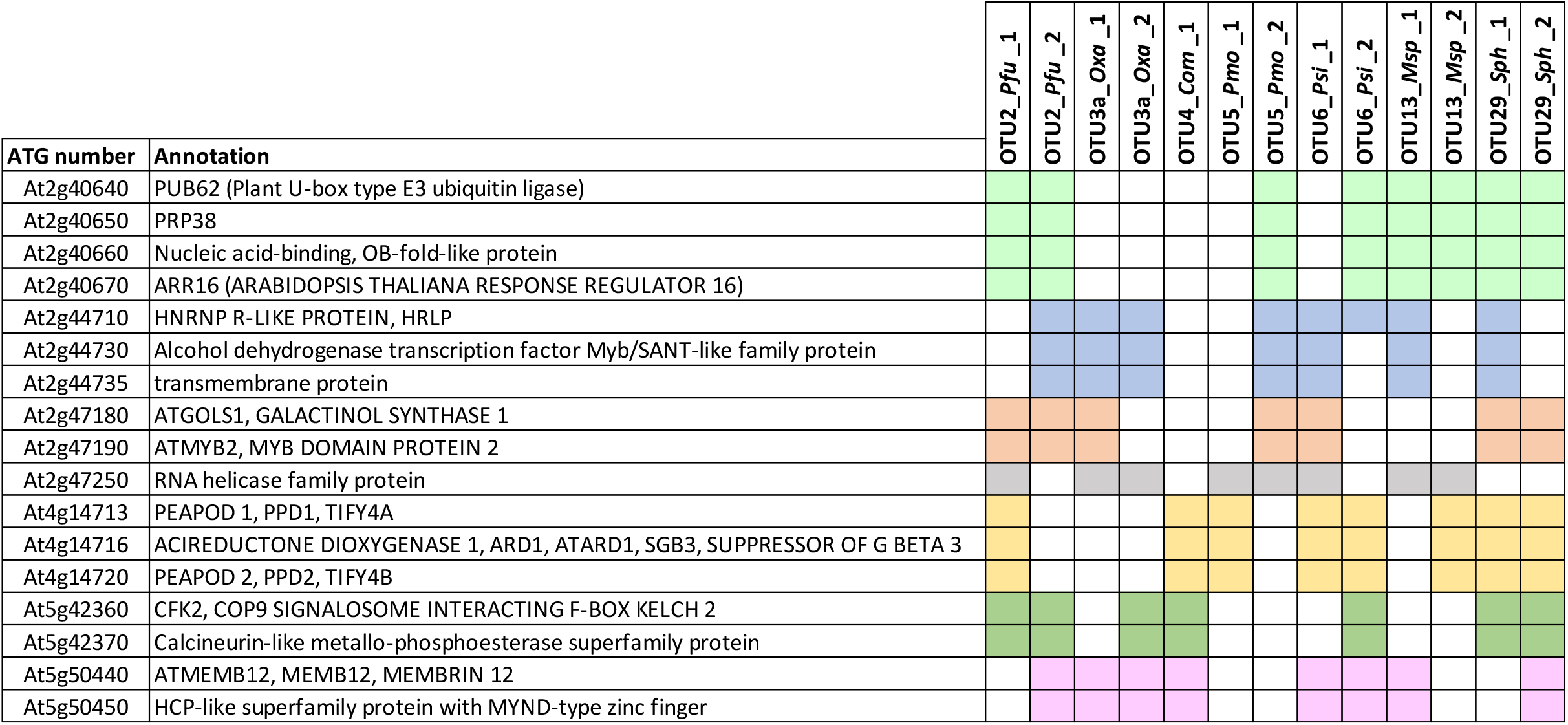
List of pleiotropic candidate genes associated with more than six bacterial strains but not detected in the mock treatment. Colored squares indicate the strains for which the candidate genes were identified. The different colors correspond to the seven QTLs in which the pleiotropic QTLs are located.

Interestingly, three genes have a link with plant immunity, *i*.*e*. the genes *MEMB12* (*At5g50440*) that is silenced by a microRNA during *Pseudomonas syringae* bacterial infection [74], *ARR16* (*At2g40670*) that is repressed by *Botrytis cinerea* fungal infection [75], and *TIFY4B/ PEAPOD 2* (*At4g14720*) that interacts with the begomovirus AL2 transcriptional activator protein, an inhibitor of plant basal defense [76].

Based on the lists of unique candidate genes identified for each treatment and the list of unique candidate genes identified across the 13 treatments with a bacterial strain, the second approach was to identify biological processes significantly over-represented in frequency compared to the overall class frequency in the *A. thaliana* MapMan annotation. When considering both the 14 treatments individually and the 13 treatments with a bacterial strain altogether, we identified 19 significantly enriched classes, among which five were also enriched in the mock treatment, i.e. ‘development’, ‘hormone metabolism’, ‘lipid metabolism’, ‘protein’ and ‘RNA’ (Figure 6a, Supplementary Data Set 5). Amongst the 14 over-represented classes not detected in the mock treatment, most of them were highly dependent on the identity of the bacterial strain, suggesting the involvement of diverse pathways in response to representative members of the non-pathogenic microbiota down to the intraspecific level (Figure 6a). We nonetheless identified four classes that were significantly over-represented for at least three treatments with a bacterial strain and when considering the 13 treatments with a bacterial strain altogether, *i*.*e*. ‘cell’, ‘secondary metabolism’, ‘signalling’ and ‘transport’ (Figure 6a). Interestingly, amongst the 99 ‘signalling’ genes, we identified (i) 54 kinase-related genes including 24 leucine-rich repeat (LRR) kinases, 8 cysteine-rich receptor-like kinases (CRK) and 6 MAP kinases, and (ii) 23 genes associated with calcium signalling, in particular for the two strains of *P. fungorum* (OTU2), the two strains of *Oxalobacteraceae* bacterium (OTU3) and one strain of *P. siliginis* (OTU6) (Figure 6b).

**Figure 6.**
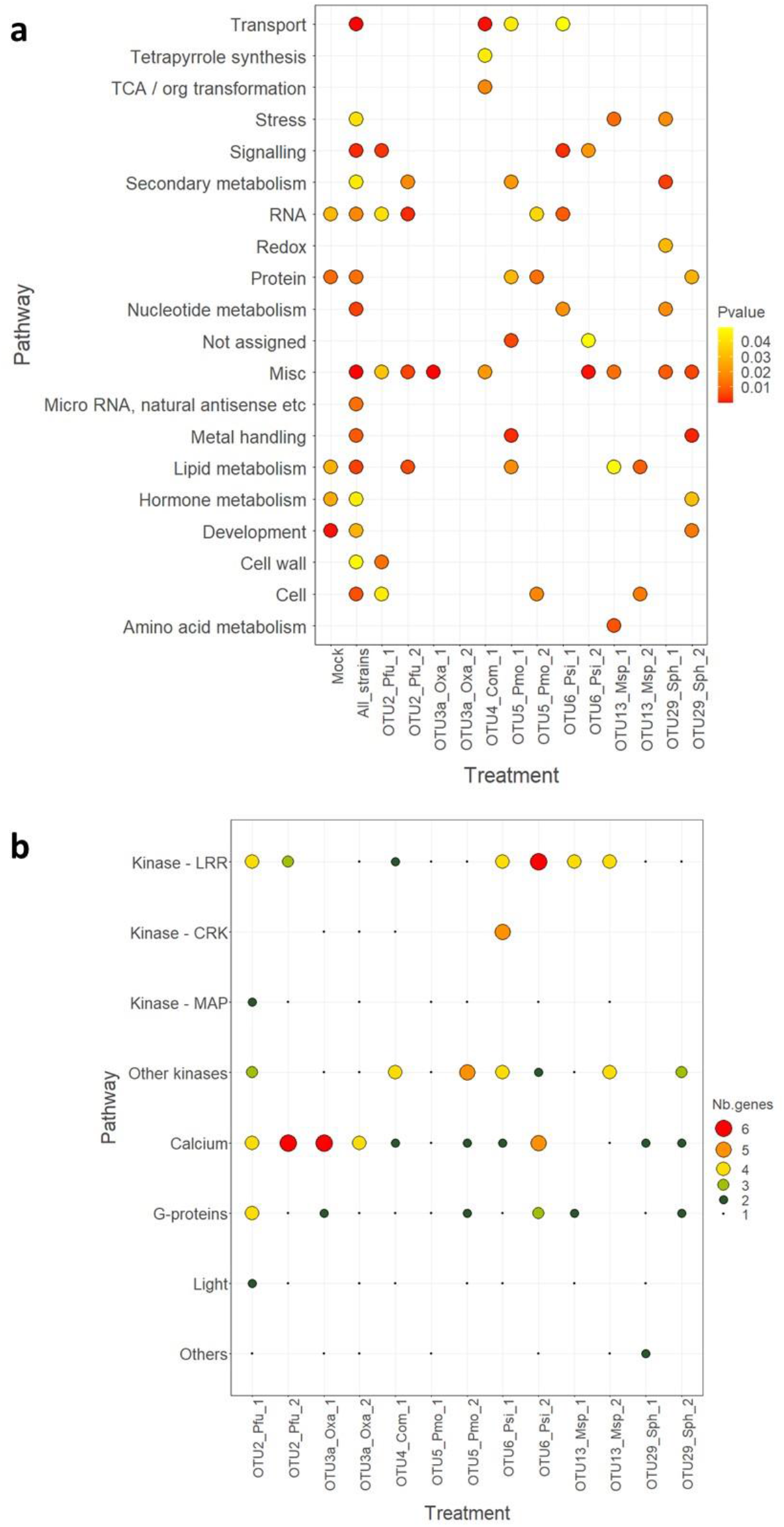
Enriched biological processes in response to the 13 bacterial strains in field conditions. **a** Enriched biological processes for the list of unique candidate genes for each of the 14 treatments and for the list of unique candidate genes from the combined 13 bacterial strains (‘All strains’), obtained with the MapMan classification superviewer tool. The color of the dots corresponds to the level of significance. **b** Number of candidate genes belonging to the different sub-categories of the enriched ‘signalling’ biological process for each treatment with a bacterial strain. LRR: leucine rich repeat, CRK: cystein-rich receptor-like kinase, MAP: mitogen-activated protein.

## DISCUSSION

### Extensive genetic variation within a local set of *A. thaliana* accessions in response to non-pathogenic leaf bacteria at the species and strain levels

Extensive genetic variation was previously observed in two worldwide collections of *A. thaliana*, each challenged at the root level in *in vitro* conditions with a single PGPB strain isolated on another plant species than *A. thaliana, i*.*e*. the strain *Pseudomonas simiae* WCS417r isolated from the rhizosphere of wheat [47] and the strain *Bacillus pumilus* TUAT-1 isolated from rice roots [49]. In this study, in line with the need to bring evolutionary and ecological functional genomics from the lab to the wild [23, 77–79], the ecological realism of plant-microbiota interactions was increased by phenotyping in field conditions, the rosette growth response of *A. thaliana* accessions collected south-west of France to non-pathogenic bacterial strains isolated from the leaf compartment of *A. thaliana* in the same geographical region.

In agreement with previous observations with bacterial pathogens [15, 31, 80], the extent of genetic variation of response to non-pathogenic bacterial strains was more dependent on the identity of the bacterial strain than the identity of the bacterial species. In addition, the presence of crossing reaction norms indicates that declaring a strain as a PGPB is highly dependent on the host genotype tested. Whether the genotype-dependent plant-growth promoting effect of a particular strain on aboveground vegetative growth is also observed at the below-ground level would deserve investigation, for instance by estimating root growth and root/shoot biomass ratios [81, 82].

Interestingly, while genetic variation in response to bacterial strains was observed within few days after inoculation in field conditions, such a genetic variation was mainly observed after several weeks in *in vitro* conditions [53]. Since the bacterial strains used in this study have been isolated from complex microbiota they used to interact and/or coevolve with in the native habitats of *A. thaliana*. Hence, the effect of bacterial strains may require the presence of additional microbiota members in the plant, a prerequisite not achieved in germ-free plants in *in vitro* conditions [53].

### A complex and highly flexible genetic architecture underlies adaptive plant-microbiota interactions

So far, the five GWAS [83–86] and the single genome-environment association study (GEAS) (Roux et al. 2022) conducted on the leaf compartment and using bacterial community descriptors as phenotypic traits, revealed a polygenic architecture controlling microbiota assembly, which is in line with the small percentage of variance explained by the phenotyping of individual mutant lines [87]. In agreement with those association genetic studies and the two GWAS conducted on *A. thaliana* in response to a PGPB strain [47, 49], we identified a complex genetic architecture for the response of *A. thaliana* to 13 non-pathogenic bacterial strains. In addition, this polygenic architecture was highly flexible among the 13 bacterial strains, with the detection of a few number of highly pleiotropic QTLs. Similar results were observed in recent GWAS conducted both in crops and wild species in response to experimental inoculation with individual pathogenic bacterial strains [88]. For instance, challenging 130 natural accessions of *A. thaliana* with 22 strains of the bacterial pathogen *Xanthomonas arboricola* revealed a clear host-strain specificity in quantitative disease resistance [80]. The complex genetic interactions observed between *A. thaliana* and the main members of its leaf microbiota should maintain high levels of diversity at the candidate genes, which in turn should result in complex co-evolutionary dynamics [16].

Beyond the question of the effects of strong genotype-by-genotype (GxG) interactions on the nature and strength of footprints of natural selection on the genome of *A. thaliana*, whether the genetic architecture underlying the response of *A. thaliana* to co-inoculation corresponds to the sum of QTLs that are specific to the response to mono-inoculations and/or to the emergence of new QTLs, remains on an open question in the research area of plant-microbe interactions. Experimental studies on plant-plant interactions demonstrated that the genetic architecture of the response of *A. thaliana* in a plurispecific neighborhood was not predictable from the genetic architecture of the response of *A. thaliana* in the corresponding bispecific neighborhoods [56, 57]

### Plant innate immunity is a significant source of natural genetic variation in plant-microbiota interactions

The candidate genes underlying the most pleiotropic QTLs have functions mainly related to plant development and/or stresses (biotic or abiotic stresses). Nevertheless, a more global approach identified four biological classes that were significantly and specifically over-represented for at least three bacterial strains but not with the mock, *i*.*e*. ‘cell’, ‘secondary metabolism’, ‘signalling’ and ‘transport’. These four classes were also over-represented in a GEAS performed on 163 natural populations of *A. thaliana* located south-west of France (including the 54 populations considered in this study) (Roux et al. 2022) and characterized *in situ* for bacterial communities in the leaf and root compartments using a metabarcoding approach [38], thereby strengthening the importance of these four classes in mediating host response to the 13 bacterial strains tested here.

Strikingly, we found a clear enrichment for signalling genes underlying QTLs in response to the 13 bacterial strains tested in this study. Signalling genes have been extensively described as being involved in plant-microbe interactions. Of particular note, we identified 8 genes belonging to the CRK family, which represents one of the largest group of RLKs with 44 members in *A. thaliana* [89, 90]. Some CRKs are involved in the regulation of plant developmental processes, while others are involved in stress and pathogen response [89]. Interestingly, by assessing host transcriptional and metabolic adaptations to 39 bacterial strains in the leaf compartment of *A. thaliana*, a core set of 24 genes consistently induced by the presence of most strains was identified and thereby referred as a molecular process called general non-self-response (GNSR) [91]. Importantly, one gene of this core set (*CRK6*) was also identified as a candidate genes in our GWAs, reinforcing the importance of CRKs in plant-microbiota interactions.

Another interesting result is that while few classical *R* genes involved in specific recognition of microbial effectors have been identified in this study, we highlighted many candidate genes related to pattern-triggered immunity (PTI), including receptor-like kinases (RLKs) and receptor-like proteins (RLPs). PTI relies on the perception of specific molecular patterns such as microbe-or pathogen-associated molecular patterns (MAMPs/PAMPs), or self-molecules (damage-associated molecular patterns, DAMPs) [92]. In particular, we identified a main actor of PTI as a candidate gene, the *FLS2* gene in response to the two strains of OTU6 and one strain of OUT 13 (Supplementary Data Set 6), the best-characterized pattern-recognition receptor (PRR) gene, encoding an LRR-RLK protein that acts as a receptor for flg22 bacterial PAMP [92, 93]. Moreover, it was previously shown that CRK6 and CRK36 are part of the PRR FLS2 protein complex, modulating PTI response through an association with FLS2 [94]. Two recent works dissected the interplay between FLS2 and numerous flg22 variants, studying how *A. thaliana* association with different evolved flg22 variants from bacterial microbiota differentially fine-tune the balance between bacterial motility and defense activation [95, 96]. PTI response is also characterized by the production of reactive oxygen species (ROS) and by the activation of the mitogen-activated protein kinases (MAPKs) cascade [97]. In our study, we identified four NADPH oxidase *RBOH* genes, among them *RBOHD* that is required for microbiota homeostasis in leaves [98]. Another candidate gene is *MPK4*, a main actor of PTI signalling (Bazin et al., 2020). Even if few mutant lines related to signalling and PTI have been tested for their effect on microbiota assembly [87, 98, 99], our results strengthen the need for a deeper investigation of some of our most promising candidate genes in relationship with the 13 strains used in this study. Importantly, the *de-novo* whole-genome sequence of the 13 strains tested in this study have been recently obtained with long-read sequencing technology [53]. Comparative genomics, and notably for their PAMP sequences (*i*.*e*. flagelline, EF-TU), may bring very informative data on their potential recognition by the plant, thereby making a link between plant-microbiota recognition and plant innate immunity. This is directly in line with a recent study that nicely shows how root commensal bacteria modulate host susceptibility to pathogens by either eliciting or dampening PTI responses [100].

## ACKNOWLEDGMENTS

We are grateful to the members of the ECOGEN tem for their assistance during the sowing and thinning. D.R.S. was funded by a PhD fellowship from CONACYT. R.D. was funded by a grant from the French Ministry of National Education and Research. This project has received funding from the European Research Council (ERC) under the European Union’s Horizon 2020 research and innovation programme (grant agreement No 951444 –PATHOCOM). This study was performed at the LIPME belonging to the Laboratoire d’Excellence (LABEX) entitled TULIP (ANR-10-LABX-41).

## DATA AVAILABILITY STATEMENT

Raw phenotypic data are available in Supplementary Data Set 1.

## SUPPLEMENTARY FIGURES AND TABLES

**Supplementary Figure S1**. Experimental design of the field experiment.

**Supplementary Figure S2**. Phenotyping of three traits related to vegetative growth by imaging (AREA, PERIMETER and DIAMETER).

**Supplementary Figure S3**. Phenotypic variation of the response to the 13 bacterial strains in field conditions.

**Supplementary Figure S4**. Box-plots illustrating the range of genetic correlations between each treatment with a bacterial strain and the remaining 13 treatments for the traits ‘perimeter-5dai’, ‘perimeter-9dai’, ‘diameter-5dai’, ‘diameter-9dai’ and ‘RGR-9dai-5dai’.

**Supplementary Figure S5**. UpSet plots illustrating the flexibility of genetic architecture among the 13 treatments with bacteria strains for the traits ‘area-5dai’, ‘perimeter-5dai’, ‘perimeter-9dai’, ‘diameter-5dai’, ‘diameter-9dai’, ‘RGR-5dai-1dbi’, ‘RGR-9dai-5dai’ and ‘RGR-9dai-1dai’.

**Supplementary Table S1**. Names and GPS coordinates (expressed in degrees) of the 54 populations used in this study.

**Supplementary Table S2**. Homogeneity of plant growth across the field trial and presence of genetic variation for the three resource acquisition traits measured on the plants before inoculation.

**Supplementary Table S3**. Genetic variation of nine traits related to resource acquisition among the 162 accessions nested within 54 populations of *A. thaliana* for each of the 14 treatments.

## SUPPLEMENTARY DATA SETS

**Supplementary Data Set 1**. Raw data for the 12 phenotypic traits scored on 14,580 plants in a field experiment conducted at INRAE Toulouse (France).

**Supplementary Data Set 2**. Genotypic values of the 54 natural populations of *A. thaliana* for the nine traits ‘area-5dai’, ‘area-9dai’, ‘perimeter-5dai’, ‘perimeter-9dai’, ‘diameter-5dai’, ‘diameter-9dai’, ‘RGR-5dai-1dbi’, ‘RGR-9dai-5dai’ and ‘RGR-9dai-1dai’ for each of the 14 treatments.

**Supplementary Data Set 3**. Genetic architecture of the 126 ‘phenotypic trait * treatment’ combinations.

**Supplementary Data Set 4**. List of all candidate genes identified for each of the 126 ‘phenotypic trait * treatment’ combinations.

**Supplementary Data Set 5**. List of unique candidate genes identified for each trait of each treatment with a bacterial strain.

**Supplementary Data Set 6**. List of the 1,962 candidate genes unique to the treatments with a bacterial strain. The pleiotropic level corresponds to the number of treatments with a bacterial strain for which the candidate gene was detected by GWA mapping.

