## Supplementary Information for "The genetic architecture of *Arabidopsis thaliana* in response to native non-pathogenic leaf bacterial species revealed by GWA mapping in field conditions"

**This file includes:**

Supplementary Text

Supplementary Figures 1-5

Supplementary Tables 1-4

### SUPPLEMENTARY TEXT

#### 1. Reducing maternal effects among the seed lots of the 162 accessions of *A. thaliana* used in this study

Differences in the maternal effects among the 162 seed lots (54 populations * 3 accessions) were reduced by growing one plant of each accession for one generation with the following steps. First, for each accession, several seeds were sown on October 1st 2016 in one 7 x 7 x 6 cm plastic pot (Soparco®) filled with damp standard culture soil (PROVEEN MOTTE 20, Soprimex®). Secondly, seeds were stratified at 4°C for four days in order to break primary dormancy and promote germination. Thirdly, pots were transferred to greenhouse conditions (22°C, 16h photoperiod) on November 4th 2016. Fourthly, seedlings were thinned to one on November 25th 2016. Fifthly, pots were placed on a field station at the INRAE campus of Auzeville (France) on December 5th 2021 in order to promote flowering through natural vernalization. Sixthly, at the onset of flowering, plants were transferred in a greenhouse mimicking the outdoor conditions (no additional light and heating) but protecting plants from rainfall. Finally, plants were separated from each other by aratubes (https://www.arasystem.com/asn003-aratubes-360) in order to avoid cross-pollination among accessions. All seed lots were harvested during a 3-week period from late April to early May 2017 and were then stored at 4°C until the set up of the phenotyping experiment.

#### 2. Phenotyping vegetative growth related traits by imaging

Three traits related to resource acquisition were measured by a non-destructive approach, by imaging each tray in the field using a digital camera (Samsung S6) in a custom box, hereafter named Photobox (Figures S2a and S2b) at 1 dbi, 5 dai and 9 dai to encompass the period of exponential growth. The three traits correspond to the projected rosette surface area (AREA), the projected rosette perimeter (PERIMETER) and the maximal projected rosette diameter (DIAMETER). To measure these three traits, we developed a color-detection based program in Python language (Supplementary text 3), with each image of a tray submitted to a seven-step treatment (Figure S2c). In the first step, the program rescales the picture of each tray and divides it in 54 individual pictures (*i.e.* one picture per well). In the second step, using the "cv2" package, the program applies two filters to each individual picture, the first one corresponding to a HSV (hue, saturation, value) color space filter and the second one corresponding to a RGB (red, green, blue) filter. In the third step, after these two color based filters, each non-white area is retrieved depending on the number of pixels implied and their location on the picture. In the fourth step, using the "mahotas" package, the program deletes all the residual non-white areas that do not correspond to plant tissues. In the fifth step, the program rescale the image around the biggest object in the picture, assuming that the largest green part of the well corresponds to the plant (it is manually verified in the next step). In the sixth step, several pictures are saved:

- The whole tray with the largest object in each well surrounded by a red square: it allows checking directly in a given tray potential problems that occur during the measurement, and thereby removing the concerned plants.
- The pictures of each well with all the pixels that exceed the threshold and one more time, the largest object surrounded by a red square.
- The plants on a white background are used by the "cv2" package to measure the parameters of interest (AREA, PERIMETER and DIAMETER).

Finally, the phenotypic data are exported in an excel file, using the package "Pandas".

In order to estimate the accuracy of the program, the maximal rosette diameter of 288 random plants (*i.e.* one tray per block) have been manually measured to the nearest millimeter at 9 dai. The correlation coefficient of Pearson between the values obtained with the program and the values estimated manually was 0.82, thereby demonstrating the great reliability of the program.

Based on this program, nine traits were therefore measured: AREA measured at 1dbi (**area-1dbi**), 5 dai (**area-5dai**) and 9 dai (**area-9dai**); PERIMETER measured at 1 dbi (**perimeter-1dbi**), 5 dai (**perimeter-5dai**) and 9 dai (**perimeter-9dai**); DIAMETER measured at 1 dbi (**diameter-1dbi**), 5 dai (**diameter-5dai**) and 9 dai (**diameter-9dai**). To estimate plant growth relative to size, the relative growth rate (RGR) of each plant was estimated based on the trait AREA:

$$\textbf{RGR-5dai-1dbi} = \frac{\ln(area-5dai)-\ln(area-1dbi)}{6\ days};$$

$$\textbf{RGR-9dai-5dai} = \frac{\ln(area-9dai)-\ln(area-5dai)}{4\ days};$$

$$\textbf{RGR-9dai-1dbi} = \frac{\ln(area-9dai)-\ln(area-1dbi)}{10\ days}$$

While the traits 'area-1dbi', 'perimeter-1dbi' and 'diameter-1dbi' allowed testing for the homogeneity of plant growth across the entire field trial (see Supplementary Table S2), the remaining nine traits were used to estimate the level of genetic variation in response to the 13 bacterial strains considered in this study.

Due to (i) the absence of plants in a well, (ii) the inability of the program to detect some non-plant objects, estimates for the 12 phenotypic traits were obtained for ~93.3% plants, *i.e.* ~14,512 plants out of 15,552 wells.

**3. Python script to measure three traits related to resource acquisition (AREA, PERIMETER and DIAMETER)**

```
"""
Created on Mon Mar  7 10:25:54 2022

@author: rduflos
"""

import numpy as np
import mahotas as mh
import cv2
import os
from pathlib import Path
from itertools import chain
import pandas as pd


#import specific functions
os.chdir("P:/Priv/Remi_Duflos/Python/table_tools")
from tabletools import merge_xlsx
kernel = np.ones((2,2),np.uint8)

def Main():
    #cwd = current working directory
    cwd = r"C:/Users/rduflos/Documents/photos_terrain/19_04_2021"
    #folder with the pictures
    rootdir = cwd + "/pics/"
    #define the working directory
    os.chdir(cwd)
    p = Path(rootdir)
    #loop through files
    for file in (chain(p.glob('**/*.jpg'))):
        #and launch functions
        images = cut_plate(file,rootdir)
        extract_plant(images)
    #this function merge the xlsx tables
    merge_xlsx (cwd)

def  cut_plate(file,rootdir):
    #initiate the programme
    images=[]
    #extract the name of the input file
    Name=file.stem
    #import the image
    image = cv2.imread(rootdir+ Name+".jpg")
    #cut the image in two groups of pots
    image_plate = image[500:4500, 240:2500].copy()
    #stock the image into list
    images.append(image_plate)
    #return a list with the image list and the name of the picture
    return images,Name

def extract_plant(images):
    #initiate a list : to establish the end table
    names,mesures,diam,perim,plant= [],[],[],[],[]
    for img in images[0] :
        plant_number =0
        image_save = img.copy()
```

```
        pixel_mm2= (54*31)/(img.shape[0]*img.shape[1])
        pixel_mm= (((540/img.shape[0])+(310/img.shape[1]))/2)

        M = int(img.shape[0]/9)
        N = int(img.shape[1]/6)
        all_height = int(M*9)
        all_width = int(N*6)

        for c in range(0,all_height,M): #walk by line and colum
            for l in range(0,all_width,N):
                plant_number +=1
                image_cut = img[c:c+M, l:l+N] #cut the image around a pot
                width = int(image_cut.shape[1])
                height = int(image_cut.shape[0]) #define the height and the
width of a pot
                ## convert to hsv
                hsv = cv2.cvtColor(image_cut, cv2.COLOR_BGR2HSV)
                ## mask of green
                maskhsv = cv2.inRange(hsv, (25, 25, 25), (100, 255,255))
                ## slice the green
                imask = maskhsv>0
                masked = np.full_like(image_cut,255, np.uint8)
                masked[imask] = image_cut[imask]
                RGB = masked.copy()
                #adjust the color keep by RGB filter
                maskRGB = cv2.inRange(RGB, (20, 65, 45), (255,200,180))
                lmask = maskRGB > 0
                image_RGBcut = np.full_like(RGB,255, np.uint8)
                image_RGBcut[lmask] = RGB[lmask]
                #define a threshold for which each element is to small to
be an Arabidpsis plant
                gauss = mh.gaussian_filter(image_RGBcut, 0.125)
                gauss = (gauss> gauss.mean())
                gauss = gauss[:,:,0]
                gauss = (gauss< gauss.mean())
                labeled, arabettes  = mh.label(gauss)
                sizes = mh.labeled.labeled_size(labeled)
                too_small = np.where(sizes < 800)
                #remove the smaller elements
                labeled = mh.labeled.remove_regions(labeled, too_small)
                # convert to rgb
                for i in range(0,height):
                      for y in range(0,width):
                          if labeled[i][y] < 1 :
                              image_RGBcut[i][y] = (255,255,255)
                #remove the noise
                image_RGBcut = cv2.morphologyEx(image_RGBcut,
cv2.MORPH_OPEN, kernel)
                if np.min(image_RGBcut) == 255 :
                    #if no variation in the image, give 0 to each value
                    names.append(images[1])
                    plant.append(str(plant_number))
                    mesures.append(0)
                    perim.append(0)
                    diam.append(0)
                else :
                    # convert to grayscale
                    gray = cv2.cvtColor(image_RGBcut, cv2.COLOR_RGB2GRAY)
                    # create a binary image based on the gray scale image
                    _, binary = cv2.threshold(gray, 225, 255,
cv2.THRESH_BINARY_INV)
```

```
                    #dilate a little bit the image to join part of the
plant
                    binarycopy = binary.copy()
                    binarydilate= cv2.dilate(binarycopy,kernel,
iterations=6)
                    #this function extract all the contours of the image
                    contours, hierarchy = cv2.findContours(
                                            image = binarydilate,
                                            mode = cv2.RETR_TREE,
                                            method =
cv2.CHAIN_APPROX_SIMPLE)
                    #initialisation, stock value to extract the biggest
contour
                    size = 0
                    biggest = 0
                    #save a copy of the image for the following steps
                    img_for_shape_area =image_RGBcut.copy()
                    for i in range (len(contours)): #loop in the contours
list to extract the biggest one
                        cnt = contours[i]
                        if cnt.size > size :
                            size = cnt.size
                            biggest = cnt
                    if type(biggest) == int :
                        print("oops")
                    else :
                        x,y,w,h = cv2.boundingRect(biggest) #find the
closest rectangle according to the biggest shape

cv2.rectangle(image_RGBcut,(x,y),(x+w,y+h),(0,0,255),2) #draw the bounding
box

cv2.rectangle(image_save,(x+l,y+c),(x+w+l,y+h+c),(0,0,255),5)
                    #create a new folder if it doesn't exist
                    if not os.path.exists("plants_in_well"):
                        os.mkdir("plants_in_well")
                        print("Directory plants_in_well Created ")
                    else:
                        pass
                    #save the image

cv2.imwrite("plants_in_well/"+images[1]+".png",image_save)

                    #save the plant with the box around
                    if not os.path.exists("plant_w_box"):
                        os.mkdir("plant_w_box")
                        print("Directory plant_w_box Created ")
                    else:
                        pass
                    cv2.imwrite("plant_w_box/"+images[1]+"_"
+str(plant_number)+".png",image_RGBcut)
                    #cut the image around the plant, defined by the box
                    plant_in_bbox =
img_for_shape_area[y:(y+h),x:(x+w)].copy()

                    #save the cut image
                    if not os.path.exists("plants"):
                        os.mkdir("plants")
                        print("Directory plants Created ")
                    else:
                        pass
```

```
                    cv2.imwrite("plants/"+images[1]+"_"
+str(plant_number)+".png",plant_in_bbox)
                    # convert to grayscale
                    gray_bbox = cv2.cvtColor(plant_in_bbox,
cv2.COLOR_RGB2GRAY)
                    # create a binary thresholded image based on the
grayscale
                    _, binary_bbox = cv2.threshold(gray_bbox, 225, 255,
cv2.THRESH_BINARY_INV)

                    binarydilate_bbox= cv2.dilate(binary_bbox,kernel,
iterations=6)

                    if not os.path.exists("binary"):
                        os.mkdir("binary")
                        print("Directory binary Created ")
                    else:
                        pass
                    cv2.imwrite("binary/"+images[1]+"_"
+str(plant_number)+"_binary.png",binarydilate_bbox)

                    #extract the contours of the new image
                    contour_plant,_ = cv2.findContours(
                                            image = binarydilate_bbox,
                                            mode = cv2.RETR_TREE,
                                            method =
cv2.CHAIN_APPROX_SIMPLE)
                    #initialisation, stock value to extract the biggest
contour
                    size_bbox = 0
                    biggest_bbox = 0

                    #list to save the sum of all the areas
                    shape_list =[]

                    for i_bbox in range (len(contour_plant)):
                        cnt_bbox = contour_plant[i_bbox]
                        #stock the areas of the shapes
                        shape_list.append(cv2.contourArea(cnt_bbox))
                        if cnt_bbox.size > size_bbox :
                            size_bbox = cnt_bbox.size
                            biggest_bbox = cnt_bbox
                    #mesuring the "area" of the visible elements
                    nzCount = cv2.countNonZero(binary_bbox)
                    a = nzCount*pixel_mm2
                    mesures.append(a)
                    #mesuring the perimeter of the visible elements
                    perimeter = (cv2.arcLength(biggest_bbox,True))*pixel_mm
                    perim.append(perimeter)
                    #mesuring the diameter of the visible elements
                    _,radius = cv2.minEnclosingCircle(biggest_bbox)
                    diameter = 2*int(radius)*pixel_mm
                    diam.append(diameter)
                    names.append(images[1])
                    plant.append(str(plant_number))
    #save into a dataframe
    df = pd.DataFrame({"name":names,"plant":plant,
"area":mesures,"perimeter":perim,"diameter":diam})
    df.to_excel(str(images[1])+".xlsx")
if "__INIT__" == "__INIT__":
    Main()
```

### SUPPLEMENTARY FIGURES

**Supplementary Figure S1.** Experimental design of the field experiment. **a** Picture of the field experiment taken by a drone. **b** Layout of the experimental design. The position of the two mock treatments (*i.e.* trays 4, 5 and 6 for treatment 'mock1' and trays 43, 44 and 45 for treatment 'mock2') were identical in each of the six experimental blocks. The 14 treatments with bacterial strains were randomized within each block.

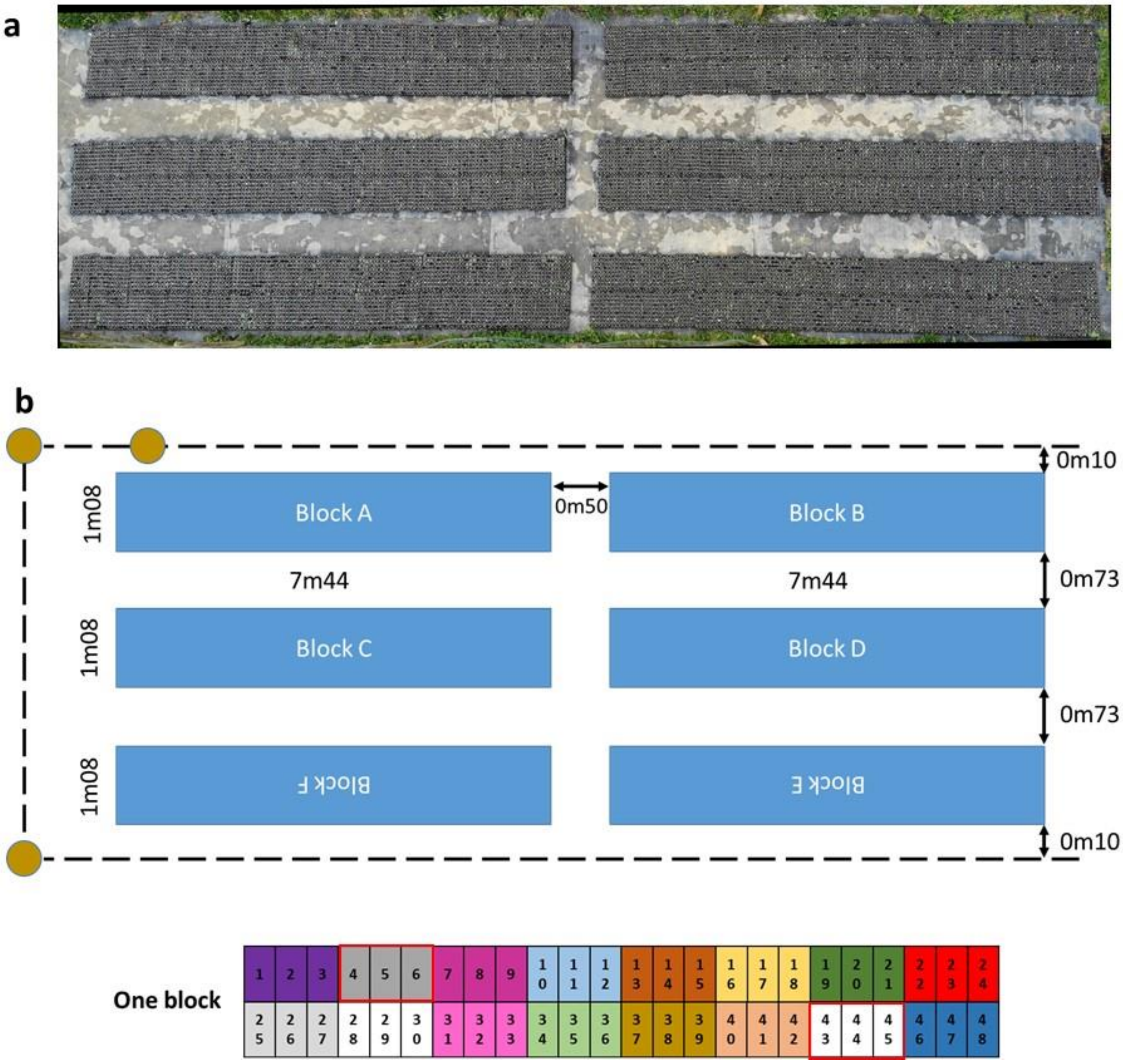

**Supplementary Figure S2.** Phenotyping of three traits related to vegetative growth by imaging (AREA, PERIMETER and DIAMETER). **a** Pictures of the photo-box used in the field. **b** Schematic side view of the photo-box. **c** Illustration of the different steps of image processing.

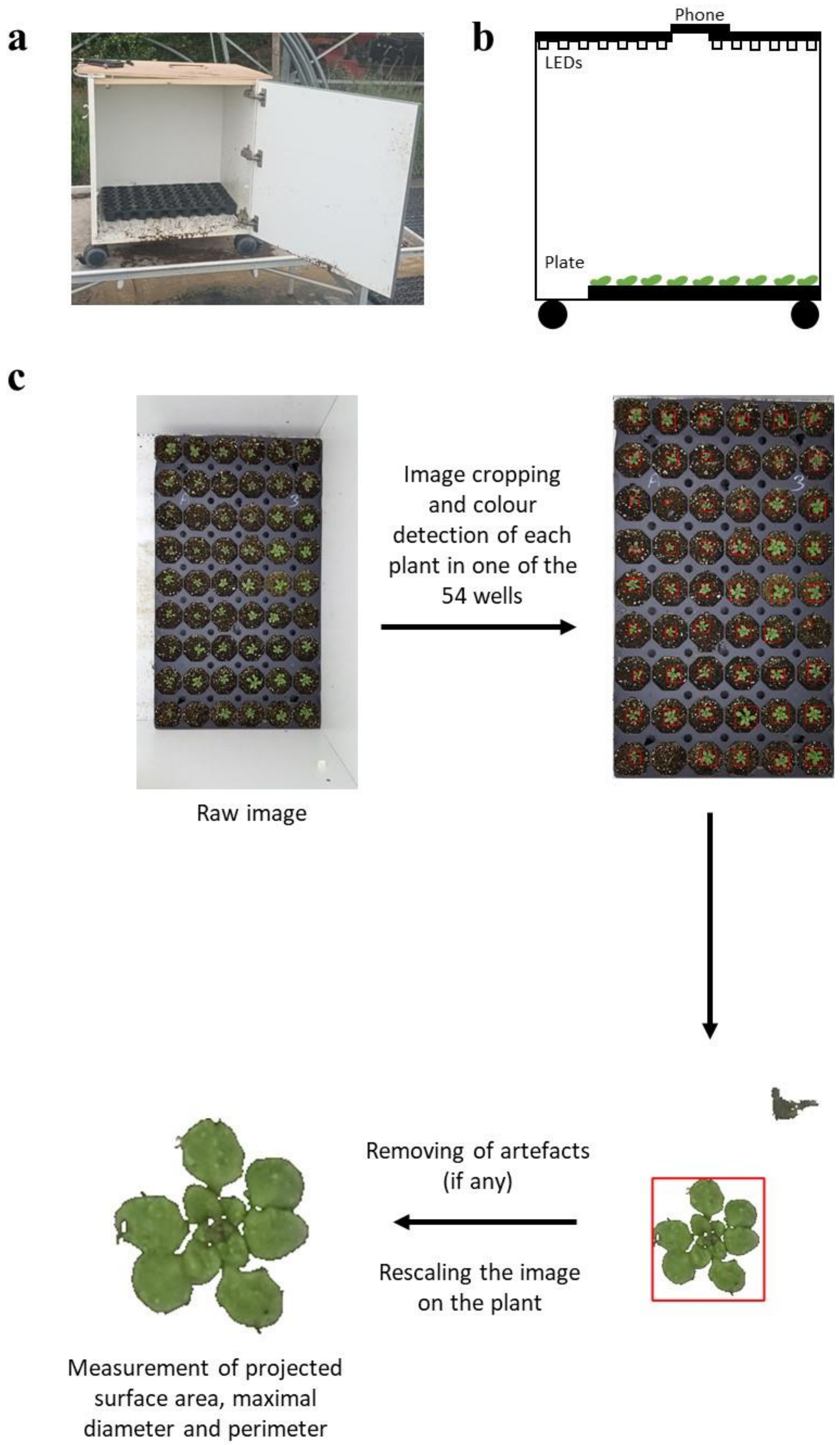

**Supplementary Figure S3.** Phenotypic variation of the response to the mock treatment and the 13 bacterial strains in field conditions. Box-plots illustrating the variation among the 14 treatments for the traits 'area-5dai', 'perimeter-5dai', 'perimeter-9dai', 'diameter-5dai', 'diameter-9dai', 'RGR-9dai-5dai' and 'RGR-9dai-1dbi'. For each treatment, each dot corresponds to the genotypic value of one of the 54 populations of *A. thaliana*. For each trait, different letters indicate different groups according to the treatments after a Ryan-Einot-Gabriel-Welsh (REGWQ) multiple-range test at $P = 0.05$. dai: days after inoculation, dbi: day before inoculation.

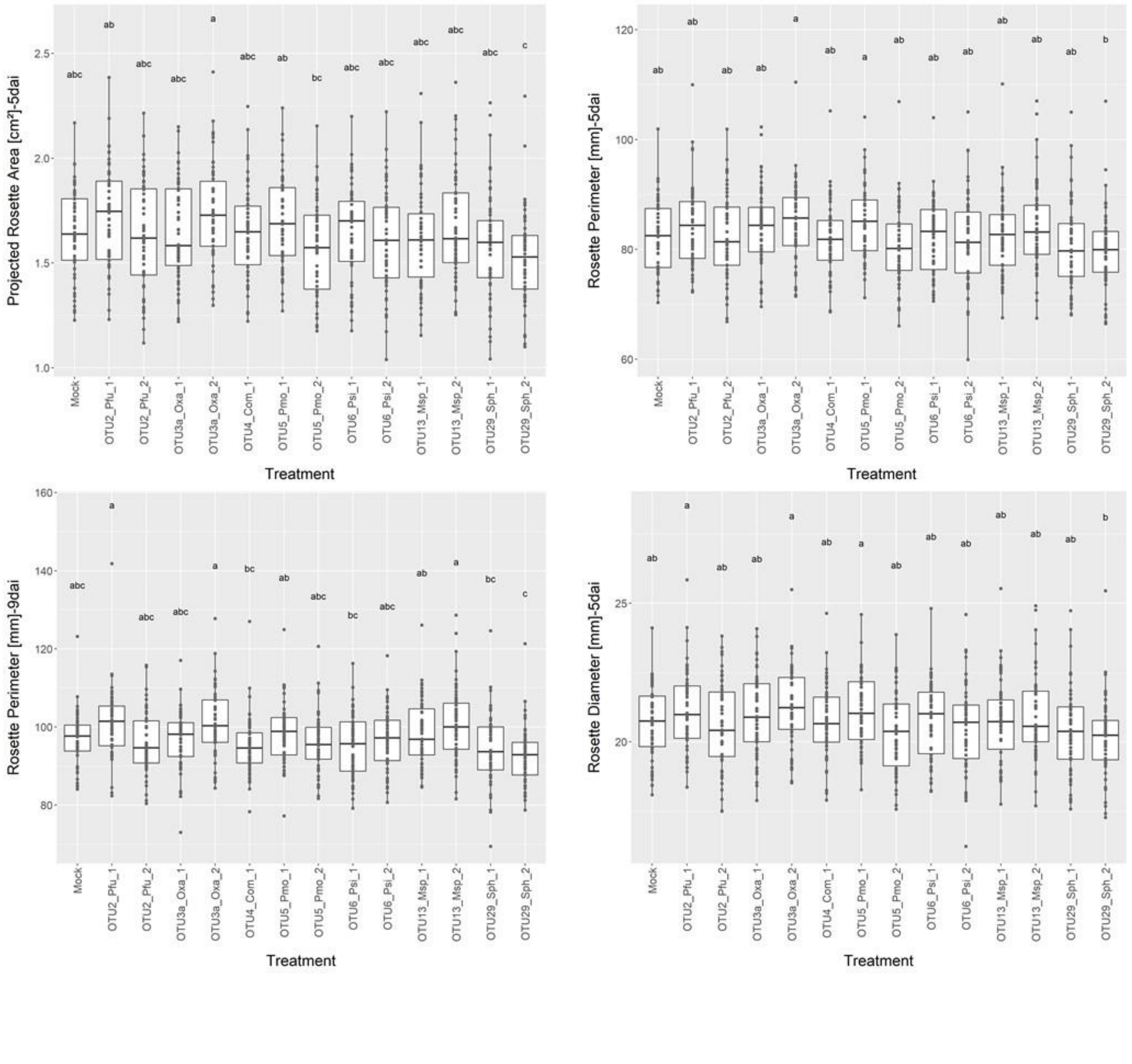

### Supplementary Figure S3 (continued)

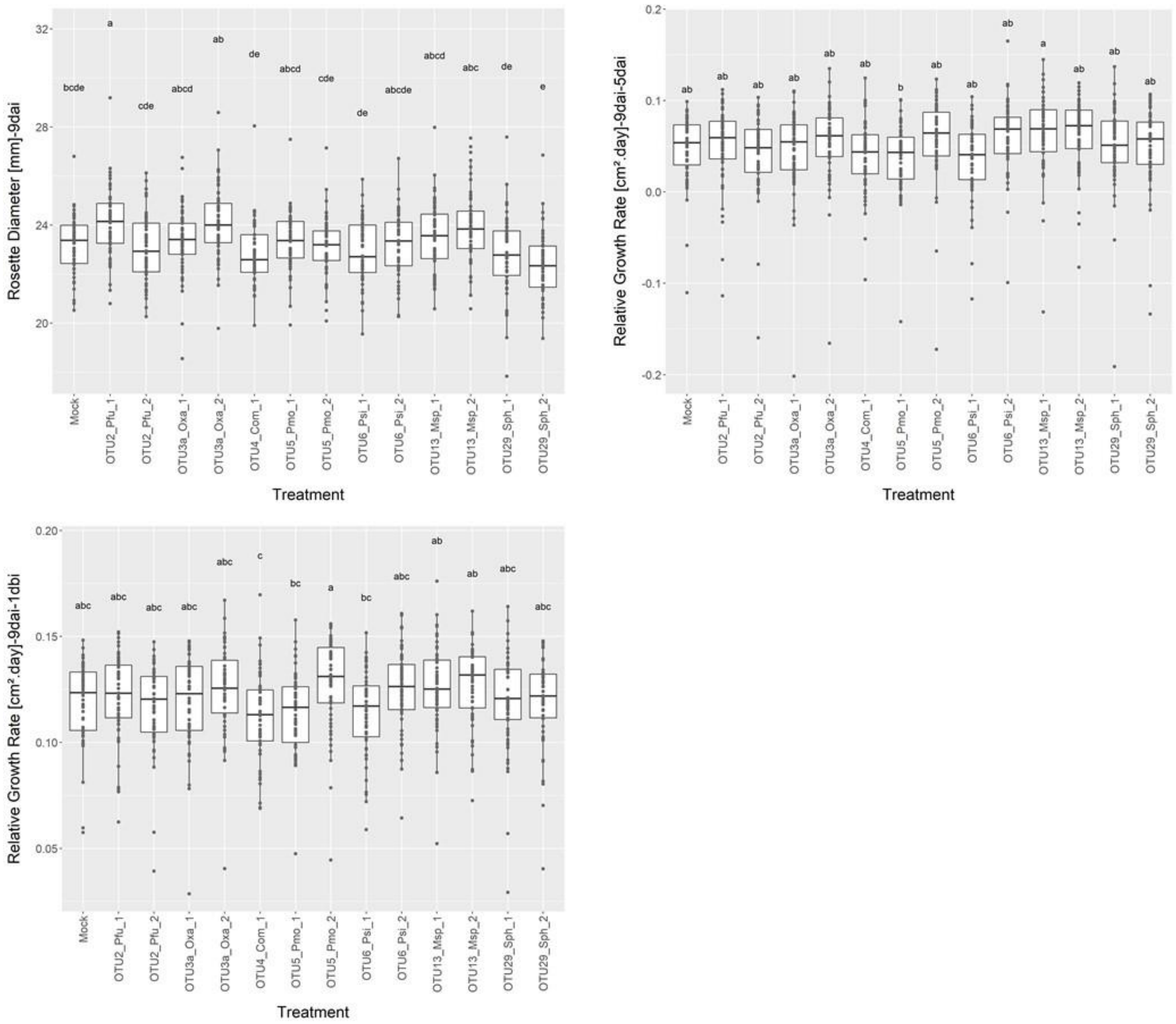

**Supplementary Figure S4**. Box-plots illustrating the range of genetic correlations between each treatment with a bacterial strain and the remaining 13 treatments for the traits ‘perimeter-5dai’, ‘perimeter-9dai’, ‘diameter-5dai’, ‘diameter-9dai’ and ‘RGR-9dai-5dai’. Red triangle: genetic correlation with the mock treatment, black dots: genetic correlations with other treatments with a bacterial strain (*ggplot2* library implemented in the R environment). dai: days after inoculation. Treatments are ranked according to their mean genetic correlation with other treatments.

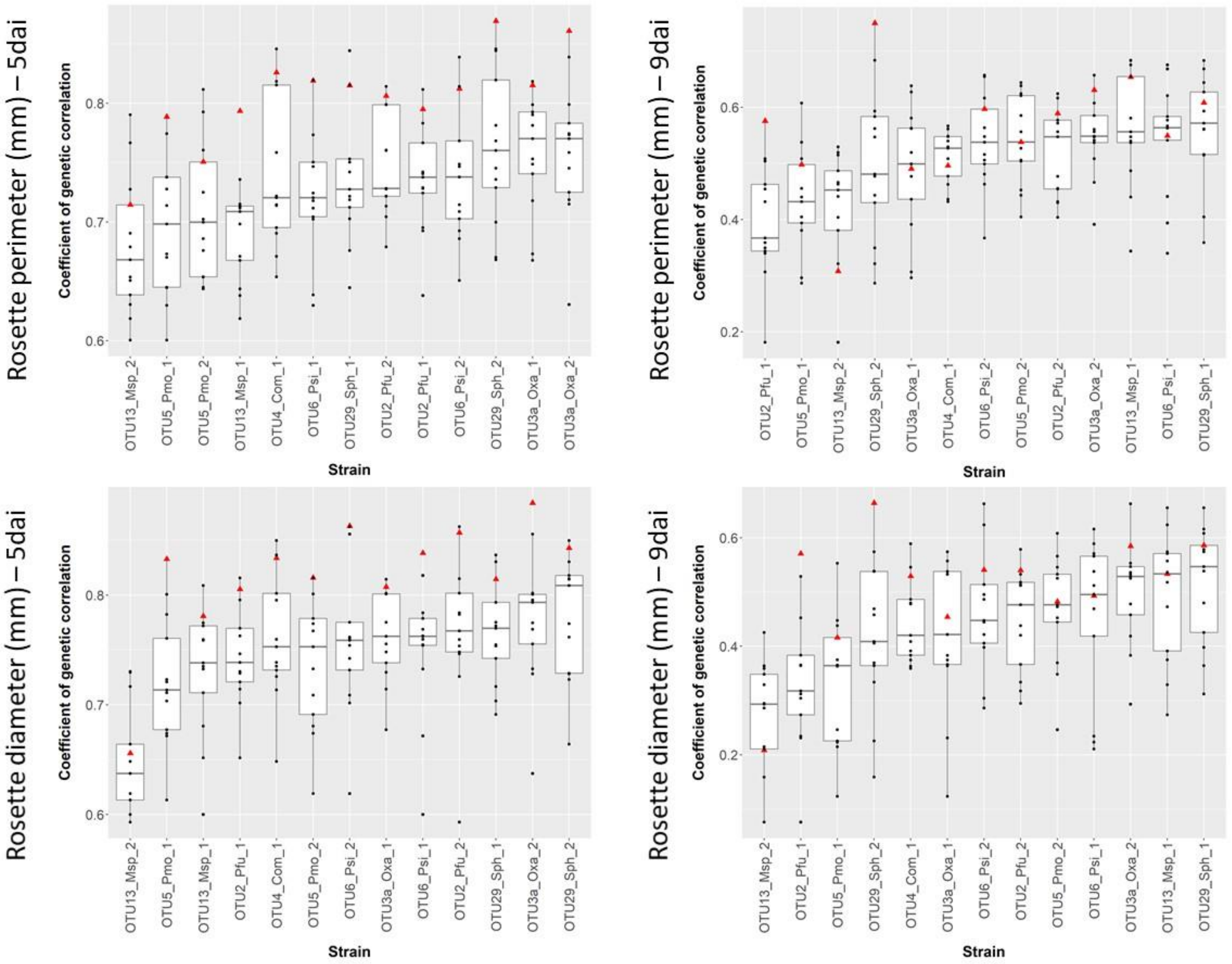

**Supplementary Figure S4 (continued)**

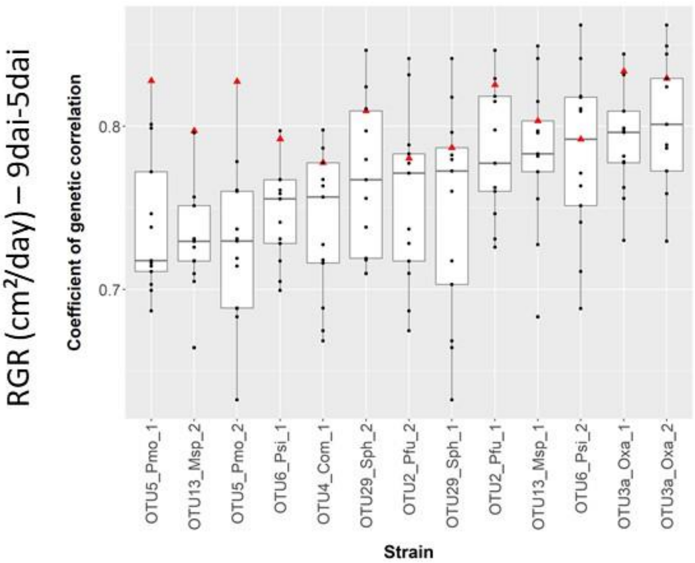

**Supplementary Figure S5**. UpSet plots illustrating the flexibility of genetic architecture among the 13 treatments with bacterial strains for the traits 'area-5dai', 'perimeter-5dai', 'perimeter-9dai', 'diameter-5dai', 'diameter-9dai', 'RGR-5dai-1dbi', 'RGR-9dai-5dai' and 'RGR-9dai-1dbi'. 'Number of genes': Total number of candidate genes underlying detected QTLs and not shared with the mock treatment. A single dot indicates the number of candidate genes specific to a given treatment. Candidate genes shared between two or more treatments are represented by a line connecting two or more dots. dai: days after inoculation; dbi: day before inoculation.

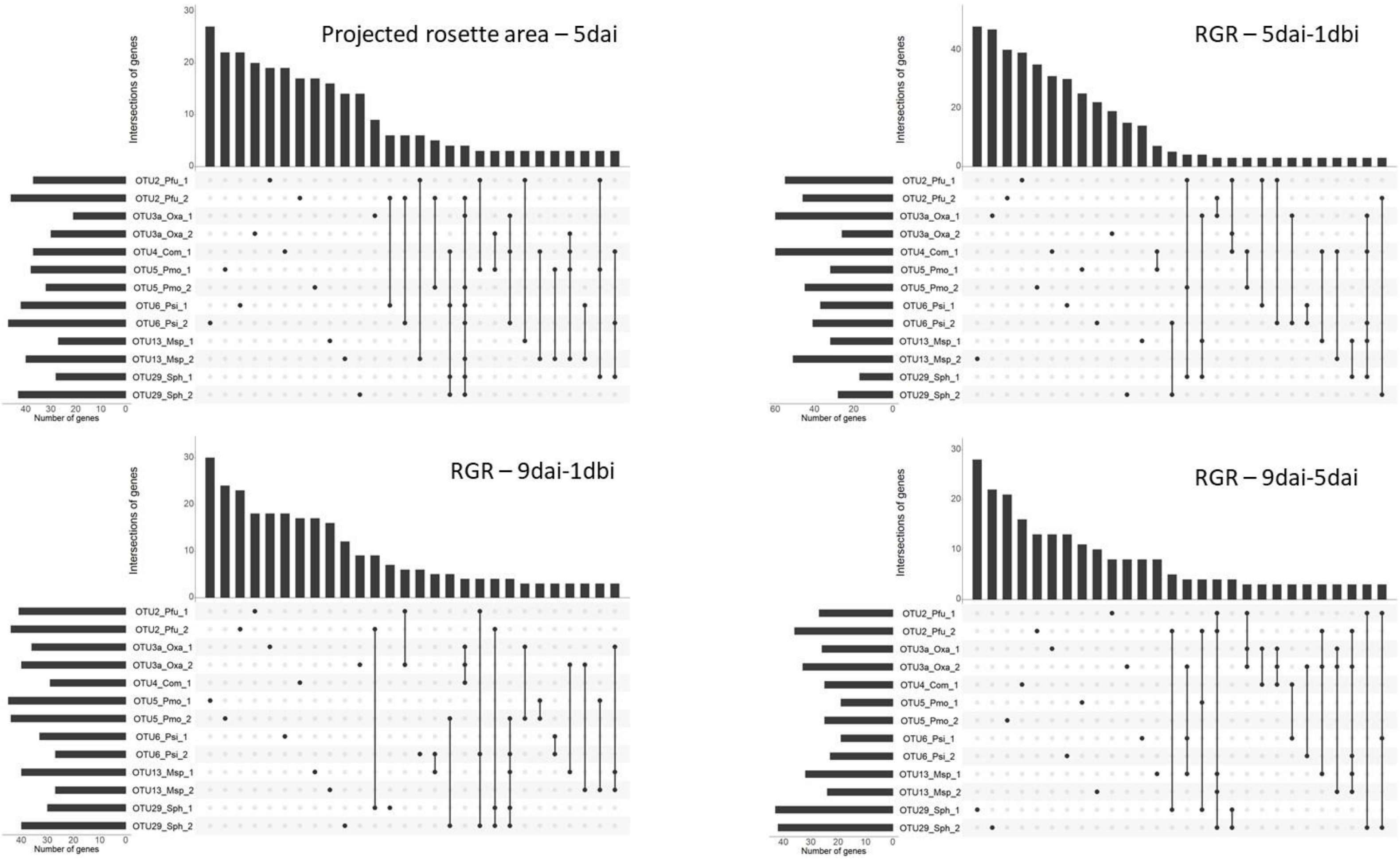

**Supplementary Figure S5 (continued)**

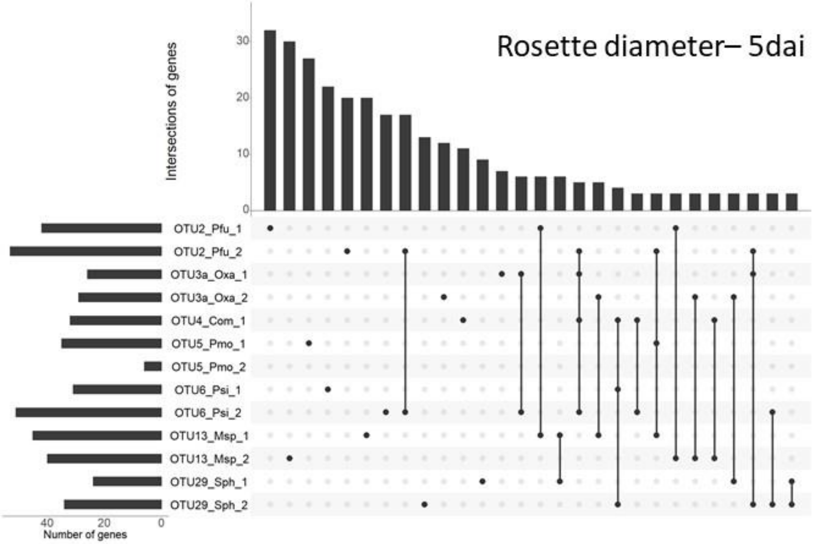


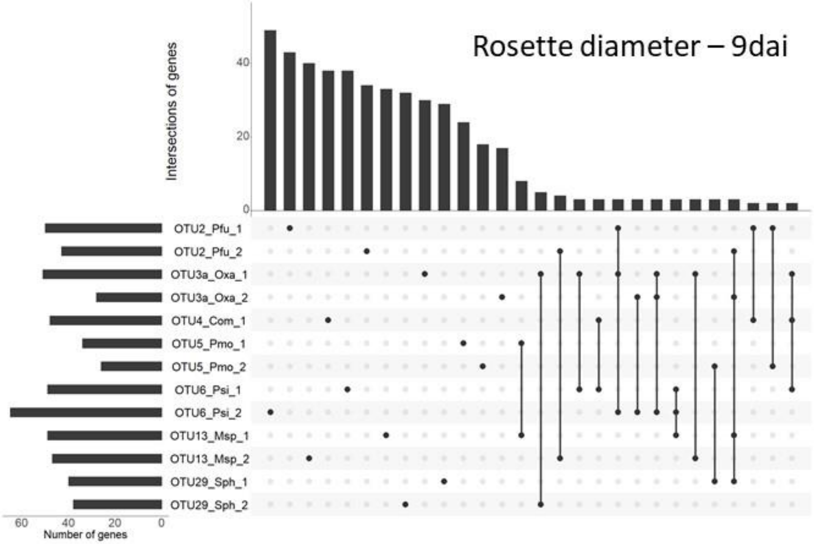


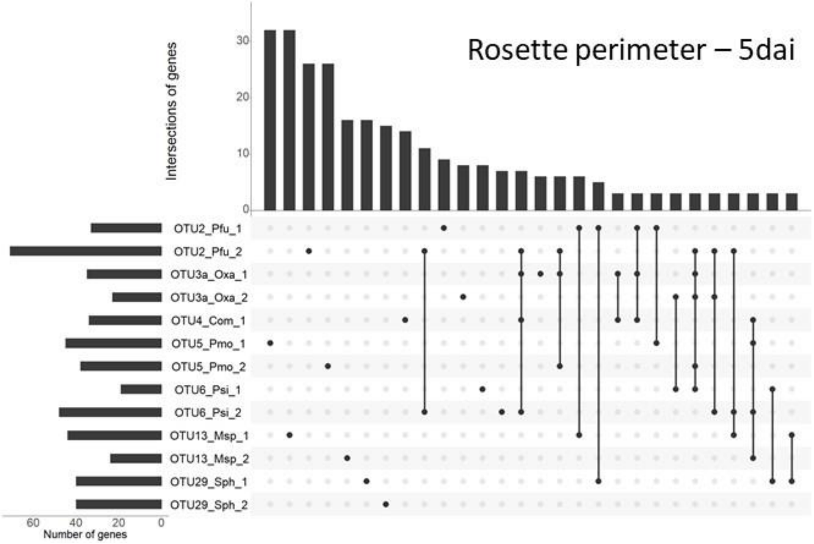


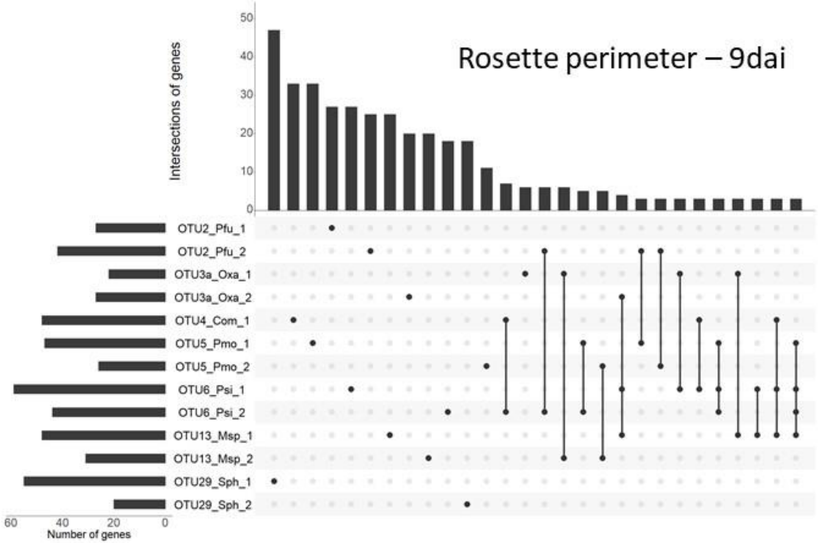

### SUPPLEMENTARY TABLES

**Supplementary Table S1.** Names and GPS coordinates (expressed in degrees) of the 54 populations used in this study.

| Population name | Locality | Latitude | Longitude |
|---|---|---|---|
| ANGE-B | Saint Angel, Salvagnac | 43.91214 | 1.656855 |
| AULO-A | Aulon | 43.190552 | 0.815774 |
| BANI-B | Banios | 43.043644 | 0.234303 |
| BARA-B | Baraqueville | 44.269727 | 2.426322 |
| BELC-C | Belcastel | 44.389212 | 2.336636 |
| CAMA-C | Camarès | 43.824878 | 2.881661 |
| CASS-A | Cassagne-Begontes | 44.17653 | 2.518164 |
| CAST-A | Castelginset | 43.698534 | 1.427856 |
| CAZA-B | Cazaux-Fréchet | 42.831484 | 0.420091 |
| CERN-A | Saint-Rome-de-Cernon | 44.01194 | 2.966488 |
| CHEI-A | Chein-dessus | 43.013708 | 0.86707 |
| CIER-A | Cier sur Luchon | 42.85332 | 0.602039 |
| CIER-B | Cier de Luchon | 42.859978 | 0.600413 |
| CLAR-A | Saint Clar-de-Rivière | 43.464776 | 1.219019 |
| CLAR-B | Saint Clar-de-Rivière | 43.465281 | 1.218577 |
| CLAR-C | Saint Clar-de-Rivière | 43.464058 | 1.21799 |
| COLO-A | Colombiès | 44.346915 | 2.340243 |
| COLO-B | Colombiès | 44.34773 | 2.339715 |
| DECA-A | Châteaude Cas (Espinas) | 44.199896 | 1.77189 |
| DIEU-A | Ville-Dieu-du-temple | 44.059797 | 1.220975 |
| ESPE-B | Esperausses | 43.693335 | 2.534582 |
| FAYA-A | Fayet | 43.8021 | 2.951709 |
| FERR-A | Ferrières | 43.657743 | 2.44371 |
| GAIL-A | Gaillac | 43.908928 | 1.900574 |
| LABA-B | Labarthe-sur-Lèze | 43.450892 | 1.40116 |
| LABA-C | Labarthe-sur-Lèze | 43.451451 | 1.39935 |
| LABA-D | Labarthe-sur-Lèze | 43.458019 | 1.381137 |
| LABAS-B | La bastide de Sérou | 43.008716 | 1.420053 |
| LACR-A | Lacraste (Montgauch) | 42.999869 | 1.075659 |
| LACR-C | Lacraste (Montgauch) | 43.000155 | 1.075624 |
| LAGR-A | Lagraulhet St Nicolas | 43.795323 | 1.073752 |
| LAMA-B | Lamasquère | 43.479745 | 1.241592 |
| LANT-C | Lanta | 43.564822 | 1.65201 |
| LUZE-B | Luzenac (Garanou) | 42.764419 | 1.753595 |
| MAZA-A | Mazamet | 43.497754 | 2.375372 |
| MERE-A | Merens-les-Vals | 42.656618 | 1.836221 |
| MERV-A | Merville | 43.720426 | 1.296824 |
| MERV-B | Merville | 43.725141 | 1.247629 |
| MONB-A | Monblanc | 43.46529 | 0.986273 |
| MONF-A | Monferran-Savès | 43.616254 | 0.972435 |
| MONT-A | Montans | 43.852212 | 1.87432 |
| MONTI-A | Montiès | 43.389383 | 0.67282 |
| MONTI-B | Montiès | 43.3839336 | 0.67257 |
| MONTM-A | Montmajou (Cier de Luchon) | 42.86156 | 0.595943 |
| MONTM-B | Montmajou (Cier de Luchon) | 42.861218 | 0.596869 |
| MOUL-A | Moularès | 44.089762 | 2.296094 |
| NAUV-B | Nauviale | 44.520418 | 2.427129 |
| NAYR-A | Le Nayrac (Cassagnes-Bégontes) | 44.161368 | 2.544711 |
| SAMA-A | Samatan | 43.494325 | 0.92391 |
| SAUB-C | Saubens | 43.475583 | 1.367589 |
| SAUR-A | Saurat | 42.889844 | 1.485209 |
| SEIS-A | Seissan | 43.487302 | 0.588798 |
| TARN-C | Villemur-sur-Tarn | 43.85328 | 1.502009 |
| VALE-A | Valence d'Albiegeois | 44.022296 | 2.403434 |

**Supplementary Table S2.** Homogeneity of plant growth across the field trial and presence of genetic variation for the three resource acquisition traits measured on the plants before inoculation. *F*: *F*-ratio, *P*: *p*-value. Bold values indicate statistically significant *p* values after correction for multiple comparisons (FDR method). dbi: day before inoculation.

| | Model terms | | | | | | | |
|---|---|---|---|---|---|---|---|---|
| | Block | | Pop | | Treatment | | Pop * Treatment | |
| Trait | *F* | *P* | *F* | *P* | *F* | *P* | *F* | *P* |
| Area_1dbi | 39.38 | ***<.0001*** | 61.05 | ***<.0001*** | 1.02 | 0.4434 | 0.79 | 1 |
| Diameter_1dbi | 41.66 | ***<.0001*** | 50.19 | ***<.0001*** | 0.74 | 0.7194 | 0.82 | 0.9997 |
| Perimeter_1dbi | 36.72 | ***<.0001*** | 45.14 | ***<.0001*** | 0.59 | 0.8517 | 0.81 | 0.9999 |

**Supplementary Table S3**. Genetic variation of nine traits related to resource acquisition among the 162 accessions nested within 54 populations of *A. thaliana* for each of the 14 treatments. *F*-ratio, *P*: *p*-value. Bold values indicate significant *p*-values after correction for multiple comparisons (FDR method). $R^2$: percentage of variance explained by model term.

| | | Area-5dai | | Area-9dai | | RGR-5dai-1dbi | | RGR-9dai-5dai | | RGR-9dai-1dbi | | Perimeter-5dai | | Perimeter-9dai | | Diameter-5dai | | Diameter-9dai | |
|---|---|---|---|---|---|---|---|---|---|---|---|---|---|---|---|---|---|---|---|
| **Treatment** | **Terms** | ***F*** | ***P*** | ***F*** | ***P*** | ***F*** | ***P*** | ***F*** | ***P*** | ***F*** | ***P*** | ***F*** | ***P*** | ***F*** | ***P*** | ***F*** | ***P*** | ***F*** | ***P*** |
| Mock | **Block** | 5.327 | ***1.32E-28*** | 2.132 | ***8.7E-06*** | 3.025 | ***1.1E-11*** | 6.213 | ***3.9E-35*** | 5.91 | ***6.2E-33*** | 6.216 | ***3.9E-35*** | 2.985 | ***2E-11*** | 5.961 | ***2.8E-33*** | 2.54 | ***2.3E-08*** |
| | **Pop** | 43.73 | ***1.65E-41*** | 23.66 | ***1.4E-22*** | 9.168 | ***1.6E-08*** | 9.449 | ***8.3E-09*** | 7.876 | ***2.8E-07*** | 50.7 | ***2.4E-47*** | 18.11 | ***3.3E-17*** | 43.13 | ***5.7E-41*** | 22.74 | ***1.1E-21*** |
| | **Accession(Pop)** | 2.304 | ***4.57E-11*** | 1.044 | 0.41 | 2.105 | ***5.7E-09*** | 2.405 | ***4.4E-12*** | 2.693 | ***2.8E-15*** | 3.411 | ***5E-24*** | 1.758 | ***1.1E-05*** | 2.763 | ***5.9E-16*** | 1.344 | ***0.019*** |
| OTU2_*Pfu* _1 | **Block** | 11.01 | ***3.79E-10*** | 1.255 | 0.117 | 1.983 | ***1.0E-04*** | 4.26 | ***7.7E-19*** | 3.206 | ***4.3E-12*** | 4.182 | ***2.2E-18*** | 2.173 | ***9.2E-06*** | 3.961 | ***5.7E-17*** | 1.552 | ***0.010*** |
| | **Pop** | 3.822 | ***4.09E-16*** | 6.032 | ***1.9E-05*** | 7.204 | ***1.5E-06*** | 9.804 | ***5.4E-09*** | 9.756 | ***6E-09*** | 12.36 | ***2.1E-11*** | 5.765 | ***3.3E-05*** | 10.75 | ***6.7E-10*** | 6.461 | ***7.5E-06*** |
| | **Accession(Pop)** | 1.746 | ***3.40E-05*** | 0.92 | 0.721 | 1.621 | ***3.2E-04*** | 1.829 | ***7.5E-06*** | 1.929 | ***1.2E-06*** | 2.493 | ***9.2E-12*** | 1.31 | ***0.035*** | 1.995 | ***3E-07*** | 1.055 | 0.382 |
| OTU2_*Pfu* _2 | **Block** | 4.681 | ***1.99E-21*** | 1.699 | ***0.002*** | 1.446 | ***0.026*** | 4.376 | ***1.8E-19*** | 3.467 | ***1.1E-13*** | 4.901 | ***8.5E-23*** | 1.945 | ***1.6E-04*** | 5.149 | ***2.3E-24*** | 1.807 | ***0.00077*** |
| | **Pop** | 41.79 | ***4.54E-37*** | 18.78 | ***3E-17*** | 21.21 | ***1.8E-19*** | 5.549 | ***5.2E-05*** | 8.731 | ***5.7E-08*** | 44.06 | ***7.6E-39*** | 21.25 | ***1.8E-19*** | 42.98 | ***4.8E-38*** | 22.67 | ***9.8E-21*** |
| | **Accession(Pop)** | 1.903 | ***2.10E-06*** | 1.106 | 0.270 | 1.847 | ***6.3E-06*** | 1.884 | ***3E-06*** | 2.085 | ***6.4E-08*** | 2.794 | ***1.6E-14*** | 1.491 | ***0.00303*** | 2.453 | ***2.8E-11*** | 1.237 | 0.084 |
| OTU3a_Oxa_1 | **Block** | 3.264 | ***1.75E-12*** | 1.693 | ***0.002*** | 1.593 | ***0.007*** | 6.141 | ***4.7E-31*** | 4.83 | ***2.2E-22*** | 3.307 | ***9.2E-13*** | 1.643 | ***0.00404*** | 3.505 | ***4.9E-14*** | 1.615 | ***0.005*** |
| | **Pop** | 14.14 | ***4.81E-13*** | 4.423 | ***0.001*** | 8.37 | ***1.2E-07*** | 33.41 | ***3.3E-30*** | 25.66 | ***2E-23*** | 14.86 | ***1.1E-13*** | 1.893 | 0.093 | 11.33 | ***1.9E-10*** | 2.444 | ***0.03299*** |
| | **Accession(Pop)** | 2.086 | ***4.83E-08*** | 1.354 | ***0.020*** | 1.343 | ***0.023*** | 2.805 | ***8.9E-15*** | 2.635 | ***4.8E-13*** | 2.49 | ***9.8E-12*** | 1.67 | ***1.4E-04*** | 2.423 | ***4E-11*** | 1.387 | ***0.013*** |
| OTU3a_Oxa_2 | **Block** | 3.81 | ***5.02E-16*** | 1.515 | ***0.01396*** | 1.722 | ***0.002*** | 5.208 | ***7.2E-25*** | 4.203 | ***1.8E-18*** | 4.243 | ***9.8E-19*** | 2.574 | ***4.1E-08*** | 3.902 | ***1.3E-16*** | 2.28 | ***2.3E-06*** |
| | **Pop** | 13.05 | ***4.81E-12*** | 11.15 | ***2.8E-10*** | 15.8 | ***1.5E-14*** | 5.975 | ***2.1E-05*** | 7.135 | ***1.7E-06*** | 12.82 | ***7.7E-12*** | 14.52 | ***2.1E-13*** | 12.5 | ***1.6E-11*** | 13.96 | ***7E-13*** |
| | **Accession(Pop)** | 2.278 | ***8.96E-10*** | 1.259 | 0.064 | 1.436 | ***0.007*** | 1.897 | ***2.1E-06*** | 1.983 | ***4E-07*** | 2.298 | ***5.8E-10*** | 1.643 | ***2.2E-04*** | 2.34 | ***2.5E-10*** | 1.492 | ***2.8E-03*** |
| OTU4_Com_1 | **Block** | 2.924 | ***2.74E-10*** | 1.553 | ***0.010*** | 2.512 | ***1E-07*** | 4.304 | ***4.1E-19*** | 4.251 | ***1.1E-18*** | 3.161 | ***8.1E-12*** | 2.084 | ***2.9E-05*** | 3.226 | ***3.2E-12*** | 1.779 | ***0.00102*** |
| | **Pop** | 20.06 | ***1.95E-18*** | 5.515 | ***5.6E-05*** | 26.05 | ***9.5E-24*** | 21.14 | ***2.1E-19*** | 19.76 | ***3.9E-18*** | 29.74 | ***4.7E-27*** | 5.57 | ***5E-05*** | 22.57 | ***1.1E-20*** | 6.648 | ***5E-06*** |
| | **Accession(Pop)** | 1.55 | ***1.10E-03*** | 0.962 | 0.629 | 1.465 | ***0.004*** | 1.972 | ***4.9E-07*** | 1.982 | ***4.4E-07*** | 1.88 | ***2.9E-06*** | 1.286 | ***0.04705*** | 1.822 | ***8.7E-06*** | 1.105 | 0.270 |
| OTU5_*Pmo* _1 | **Block** | 2.643 | ***1.53E-08*** | 0.873 | 0.727 | 1.753 | ***0.001*** | 3.455 | ***1E-13*** | 2.874 | ***5.7E-10*** | 2.904 | ***3.5E-10*** | 1.704 | ***0.00225*** | 2.773 | ***2.4E-09*** | 1.221 | 0.147 |
| | **Pop** | 31.66 | ***8.90E-29*** | 15.88 | ***1.3E-14*** | 3.152 | ***0.008*** | 26.94 | ***1.3E-24*** | 13.49 | ***1.9E-12*** | 33.09 | ***5.4E-30*** | 12.88 | ***6.8E-12*** | 28.67 | ***3.8E-26*** | 16.73 | ***2.1E-15*** |
| | **Accession(Pop)** | 1.658 | ***1.73E-04*** | 1.194 | 0.126 | 1.911 | ***1.6E-06*** | 2.12 | ***2.3E-08*** | 2.442 | ***2.8E-11*** | 1.888 | ***2.4E-06*** | 1.353 | ***0.020*** | 1.795 | ***1.4E-05*** | 1.217 | 0.102 |
| OTU5_*Pmo* _2 | **Block** | 3.438 | ***1.41E-13*** | 1.282 | 0.097 | 1.833 | ***0.001*** | 6.186 | ***3.4E-31*** | 4.66 | ***2.7E-21*** | 3.795 | ***6.9E-16*** | 1.916 | ***2.3E-04*** | 3.635 | ***7.5E-15*** | 1.493 | ***0.02*** |
| | **Pop** | 38.68 | ***1.42E-34*** | 13.19 | ***3.7E-12*** | 12.2 | ***3.1E-11*** | 14.74 | ***1.4E-13*** | 7.52 | ***7.7E-07*** | 35.87 | ***3E-32*** | 15.72 | ***1.8E-14*** | 35.95 | ***2.6E-32*** | 15.6 | ***2.2E-14*** |
| | **Accession(Pop)** | 1.773 | ***2.21E-05*** | 1.176 | 0.146 | 0.906 | 0.753 | 2.016 | ***2.1E-07*** | 1.734 | ***4.5E-05*** | 1.753 | ***3.2E-05*** | 1.266 | 0.060 | 1.729 | ***4.8E-05*** | 1.1 | 0.280 |
| OTU6_*Psi* _1 | **Block** | 3.903 | ***1.39E-16*** | 1.286 | 0.095 | 1.895 | ***0.000*** | 4.096 | ***8.9E-18*** | 3.333 | ***7.1E-13*** | 3.769 | ***9.8E-16*** | 2.065 | ***3.7E-05*** | 4.141 | ***4.7E-18*** | 1.658 | ***0.003*** |
| | **Pop** | 17.99 | ***1.51E-16*** | 13.27 | ***3.1E-12*** | 7.998 | ***2.7E-07*** | 9.634 | ***7.8E-09*** | 7.113 | ***1.8E-06*** | 15.94 | ***1.2E-14*** | 11.49 | ***1.4E-10*** | 15.9 | ***1.3E-14*** | 15.61 | ***2.2E-14*** |
| | **Accession(Pop)** | 1.935 | ***1.05E-06*** | 0.94 | 0.678 | 1.56 | ***0.001*** | 2.168 | ***9.6E-09*** | 2.219 | ***3.5E-09*** | 2.629 | ***5E-13*** | 1.43 | ***0.01*** | 2.459 | ***2.1E-11*** | 1.214 | 0.106 |
| OTU6_*Psi* _2 | **Block** | 4.22 | ***1.48E-18*** | 1.027 | 0.434 | 1.838 | ***0.001*** | 3.702 | ***2.6E-15*** | 3.217 | ***3.9E-12*** | 4.789 | ***3.4E-22*** | 1.895 | ***0.00028*** | 4.673 | ***2E-21*** | 1.42 | ***0.03259*** |
| | **Pop** | 47.98 | ***5.19E-42*** | 16.15 | ***7.4E-15*** | 9.83 | ***5.2E-09*** | 39.2 | ***5.1E-35*** | 27.75 | ***2.9E-25*** | 55.5 | ***2.4E-47*** | 23.26 | ***2.5E-21*** | 49.36 | ***4.8E-43*** | 20.59 | ***6.2E-19*** |
| | **Accession(Pop)** | 1.923 | ***1.31E-06*** | 1.183 | 0.137 | 1.85 | ***5.5E-06*** | 1.78 | ***1.9E-05*** | 2.052 | ***1E-07*** | 2.325 | ***3.4E-10*** | 1.642 | ***0.00022*** | 2.278 | ***9.1E-10*** | 1.35 | ***0.021*** |
| OTU13_Msp_1 | **Block** | 3.926 | ***1.02E-16*** | 1.506 | ***0.015*** | 1.956 | ***1.4E-04*** | 4.531 | ***1.5E-20*** | 3.648 | ***6.7E-15*** | 3.713 | ***2.2E-15*** | 2.483 | ***1.5E-07*** | 4.118 | ***6.2E-18*** | 1.966 | ***1.3E-04*** |
| | **Pop** | 36.99 | ***3.30E-33*** | 14.25 | ***3.9E-13*** | 8.008 | ***2.7E-07*** | 69.14 | ***4.5E-57*** | 43.47 | ***2.2E-38*** | 35.24 | ***9.6E-32*** | 9.236 | ***1.8E-08*** | 31.53 | ***1.3E-28*** | 13.71 | ***1.2E-12*** |
| | **Accession(Pop)** | 2.775 | ***1.65E-14*** | 1.698 | ***8.5E-05*** | 1.359 | ***0.01947*** | 2.216 | ***3.5E-09*** | 2.067 | ***7.8E-08*** | 3.078 | ***2.8E-17*** | 2.097 | ***4E-08*** | 3.223 | ***1E-18*** | 1.968 | ***5.2E-07*** |
| OTU13_Msp_2 | **Block** | 3.949 | ***7.93E-17*** | 1.707 | ***0.002*** | 1.809 | ***7.6E-04*** | 3.805 | ***6.2E-16*** | 3.183 | ***6.5E-12*** | 3.929 | ***1E-16*** | 2.225 | ***4.9E-06*** | 3.904 | ***1.4E-16*** | 1.822 | ***0.001*** |
| | **Pop** | 48.74 | ***1.49E-42*** | 14.76 | ***1.3E-13*** | 16.27 | ***6.1E-15*** | 18.19 | ***1E-16*** | 8.288 | ***1.5E-07*** | 47.38 | ***1.6E-41*** | 14.9 | ***1E-13*** | 47.46 | ***1.4E-41*** | 14.74 | ***1.4E-13*** |
| | **Accession(Pop)** | 2.422 | ***5.02E-11*** | 1.188 | 0.134 | 1.139 | 0.205884 | 2.159 | ***1.3E-08*** | 1.852 | ***5.6E-06*** | 2.661 | ***3.4E-13*** | 1.352 | ***0.02104*** | 2.613 | ***8.8E-13*** | 1.293 | ***0.045*** |
| OTU29_Sph_1 | **Block** | 4.45 | ***4.56E-20*** | 2.196 | ***6.9E-06*** | 2.586 | ***3.6E-08*** | 4.953 | ***3.4E-23*** | 4.789 | ***3.4E-22*** | 4.54 | ***1.2E-20*** | 3.001 | ***8.6E-11*** | 4.571 | ***8E-21*** | 2.62 | ***2.2E-08*** |
| | **Pop** | 49.73 | ***2.35E-43*** | 19.12 | ***1.4E-17*** | 21.21 | ***1.8E-19*** | 9.591 | ***8.4E-09*** | 7.769 | ***4.4E-07*** | 54.85 | ***4.9E-47*** | 27.63 | ***3.1E-25*** | 51.84 | ***6.8E-45*** | 24.98 | ***7.4E-23*** |
| | **Accession(Pop)** | 1.454 | ***5.02E-03*** | 0.967 | 0.627 | 1.653 | ***1.9E-04*** | 2.053 | ***9.5E-08*** | 2.339 | ***2.5E-10*** | 2.16 | ***1.1E-08*** | 1.088 | 0.304 | 1.863 | ***3.9E-06*** | 0.961 | 0.629 |
| OTU29_Sph_2 | **Block** | 3.428 | ***1.59E-13*** | 1.808 | ***0.00076*** | 1.72 | ***0.00196*** | 5.219 | ***7.2E-25*** | 4.446 | ***5.2E-20*** | 4.032 | ***2.2E-17*** | 1.935 | ***1.8E-04*** | 3.736 | ***1.6E-15*** | 1.859 | ***0.00043*** |
| | **Pop** | 31.45 | ***1.52E-28*** | 7.493 | ***8E-07*** | 17.42 | ***5E-16*** | 32.25 | ***3.3E-29*** | 27.81 | ***2.5E-25*** | 32.47 | ***2.1E-29*** | 11.94 | ***5.2E-11*** | 28.24 | ***1E-25*** | 9.874 | ***4.7E-09*** |
| | **Accession(Pop)** | 1.422 | ***8.02E-03*** | 0.874 | 0.820 | 1.964 | ***5.8E-07*** | 1.912 | ***1.6E-06*** | 2.171 | ***9.3E-09*** | 1.921 | ***1.3E-06*** | 1.097 | 0.286 | 1.643 | ***2.2E-04*** | 0.869 | 0.827 |

**Supplementary Table S4**. **Broad-sense heritability (*H*²) estimates based on 162 accessions of *A. thaliana*, for each of the 126 'phenotypic trait * treatment' combinations.** Bold values indicate significant *H*² estimates after correction for multiple comparisons (FDR method).

| Trait | Mock | OTU2_Pfu_1 | OTU2_Pfu_2 | OTU3a_Oxa_1 | OTU3a_Oxa_2 | OTU4_Com_1 | OTU5_Pmo_1 | OTU5_Pmo_2 | OTU6_Psi_1 | OTU6_Psi_2 | OTU13_Msp_1 | OTU13_Msp_2 | OTU29_Sph_1 | OTU29_Sph_2 |
|---|---|---|---|---|---|---|---|---|---|---|---|---|---|---|
| area-5dai | **0.54** | **0.58** | **0.65** | **0.59** | **0.64** | **0.49** | **0.48** | **0.58** | **0.61** | **0.63** | **0.70** | **0.66** | **0.60** | **0.53** |
| area-9dai | **0.17** | **0.03** | **0.25** | **0.32** | **0.25** | **0.13** | 0.06 | **0.19** | 0.05 | **0.09** | **0.39** | **0.26** | **0.30** | **0.17** |
| RGR-5dai-1dbi | **0.42** | **0.44** | **0.37** | **0.29** | **0.36** | **0.45** | **0.46** | **0.20** | **0.40** | **0.47** | **0.37** | **0.27** | **0.48** | **0.48** |
| RGR-9dai-5dai | **0.58** | **0.63** | **0.63** | **0.75** | **0.67** | **0.65** | **0.62** | **0.71** | **0.66** | **0.59** | **0.69** | **0.64** | **0.68** | **0.68** |
| RGR-9dai-1dbi | **0.59** | **0.58** | **0.61** | **0.71** | **0.64** | **0.65** | **0.63** | **0.64** | **0.62** | **0.59** | **0.64** | **0.58** | **0.69** | **0.67** |
| perimeter-5dai | **0.63** | **0.67** | **0.72** | **0.63** | **0.67** | **0.56** | **0.53** | **0.60** | **0.67** | **0.69** | **0.72** | **0.68** | **0.68** | **0.62** |
| perimeter-9dai | **0.37** | **0.36** | **0.40** | **0.40** | **0.49** | **0.33** | **0.31** | **0.33** | **0.38** | **0.41** | **0.56** | **0.39** | **0.44** | **0.27** |
| diameter-5dai | **0.59** | **0.61** | **0.71** | **0.64** | **0.65** | **0.55** | **0.51** | **0.58** | **0.67** | **0.68** | **0.74** | **0.68** | **0.65** | **0.58** |
| diameter-9dai | **0.27** | **0.16** | **0.32** | **0.31** | **0.44** | **0.23** | **0.16** | **0.19** | **0.24** | **0.23** | **0.50** | **0.31** | **0.36** | **0.17** |
